# Multifaceted diversity traits of crucial microbial groups in biological soil crusts promote soil multifunctionality

**DOI:** 10.1101/797191

**Authors:** Hua Li, Youxin Chen, Gongliang Yu, Federico Rossi, Da Huo, Roberto De Philippis, Xiaoli Cheng, Weibo Wang, Renhui Li

**Author notes:** WBW and RHL are corresponding authors. Author for correspondence: Dr. Hua Li (Institute of Hydrobiology, Chinese Academy of Sciences); Address: South Donghu Road 7, 430072 Wuhan, China. Department of Food, Environmental and Nutritional Sciences, University of Milan, Milano 20133, Italy.

## Abstract

Microbial diversity is one of the most important drivers on ecosystem to maintain the simultaneous performance of functions (multifunctionality, MF) under climatic oscillation. However, existing studies typically consider taxonomic richness or Shannon index at the community level in which relations between diversity and functioning are not highly consistent. To disentangle the underlying linkages in real-world ecosystems, we conducted field investigation on biological soil crusts of Tibetan Plateau and evaluated multiple diversity facets (*i.e.*, richness, evenness, and phylogeny-related trait dissimilarity) of carbon- and nitrogen-fixing functional groups (FGs). Seven crucial variables of soil functioning were also identified to calculate MF. We found that the integrated index, invoking multiple diversity components, was a stronger predictor on MF than richness. Moreover, the divergent performance of different diversity facets determined the idiosyncratic diversity effect of each FG on the MF. Namely, richness was the dominant factor for diazotrophs to maximize MF, whereas phylogenetic dissimilarity was the most important one for phototrophs. The heterogeneity among the focal FGs derived from the significant differentiation of the extent of multifunctional redundancy. Collectively, we speculated that the multifaceted diversity pattern depicts the response ability of crucial FGs by which biocrusts stabilize MF under environmental perturbation. Taken together, our results provided a perspective to bridge the gap between taxonomic and trait-based approaches for elucidating the biodiversity-ecosystem functioning relationship, and could ultimately help to boost the practices of dryland management against global change.

## 1. Introduction

The era of global changes is characterized as a period of rapid reduction in the number of species that comprise biodiversity (Cadotte, Cardinale, & Oakley, 2008; F. T. Maestre et al., 2016). A growing body of evidence highlights that biodiversity does not merely respond to environmental fluctuations but also determines ecosystem properties at various biological scales (Cardinale et al., 2012; Carey, 2016; Hector et al., 1999; Loreau et al., 2001). The potential for biodiversity loss to impair the delivery of ecosystem functions (EFs) and services has motived researchers to better understand the underlying linkages between biodiversity and ecosystem functioning (Allan et al., 2015; Cernansky, 2017; Mori, Furukawa, & Sasaki, 2013; Tilman, Wedin, & Knops, 1996; Zavaleta, Pasari, Hulvey, & Tilman, 2010). In particular, higher levels of biodiversity are found to be certainly required to maintain the simultaneous performance of multiple EFs (‘multifunctionality’, hereafter MF) (M.A. Bowker, Maestre, & Mau, 2013; Byrnes et al., 2014; Hector & Bagchi, 2007).

As one of the most complex habitats on Earth, soils contain an immense diversity of microorganisms (Bardgett & van der Putten, 2014; Wagg, Bender, Widmer, & van der Heijden, 2014; Wall, 1999). The awareness is increasing among ecologists that soil microbes play a crucial role on the stability and resilience of terrestrial ecosystems (Bastida et al., 2016; Carey, 2016; Delgado-Baquerizo et al., 2017; Yang et al., 2014). Microbial diversity in soils has attracted wide attention over the last decade (Baveye, Berthelin, & Munch, 2016). However, this mega-diverse community remains largely unknown (Bender, Wagg, & van der Heijden, 2016; Carey, 2016). We are just beginning to test how soil microbial diversity explicitly responds to external perturbation and elucidate the mechanisms by which it drives soil MF (Baveye et al., 2016; Delgado-Baquerizo, Maestre, et al., 2016; Jing et al., 2015; Mori et al., 2016). Emerging studies, mainly conducted on the metric of taxonomic diversity at the community level, suggest that microbial diversity is of key importance on driving ecosystem functioning, even simultaneously accounting for spatial and environmental drivers (Castillo-Monroy et al., 2011; Delgado-Baquerizo et al., 2017; Delgado-Baquerizo, Maestre, et al., 2016; Mori et al., 2016; Soliveres et al., 2016). Yet, the general relationship between microbial diversity and MF is not highly consistent among different taxonomic groups (Bradford et al., 2014). For instance, an observational study on soil biodiversity (richness) of Tibetan Plateau showed that, despite positive relations of bacteria and microfauna with MF, some other groups (archaea and fungi) are not related to MF significantly (Jing et al., 2015). This kind of discontinuity between groups makes it difficult to aid the practical application of knowledge about diversity-functioning relations (Bradford et al., 2014; Mori et al., 2013).

Recent studies argue that traditional taxonomic measures may not adequately capture the features of biodiversity that most correlated with ecosystem functioning (Bender et al., 2016; Cernansky, 2017; Mori et al., 2013). The alternative view is proposed to concern on trait-based measures in terms of functional groups (FGs) (Krause et al., 2014; Mori et al., 2013). An example focusing on metabolic functions of microbial communities has demonstrated that the degradation of chitin and cellulose are controlled by single FGs and their trait rather than by the diversity of the overall community (Peter et al., 2010). Given the inherently mechanistic correlation (*i.e.*, loss of any FGs will likely result in loss of some functioning) (Mori et al., 2013), relating particular functioning processes to the diversity of special functionally coherent groups is likely to facilitate our understanding on the linkages between microbial diversity and MF (Krause et al., 2014). Nevertheless, sustaining more of everything of various FGs in an unspecified manner might not in practice be rational and realizable (Bender et al., 2016). Meanwhile, it’s noteworthy that current biodiversity loss is nonrandom and probably causes crucial losses of species for special FGs that contribute more largely to MF (Selmants, Zavaleta, Pasari, & Hernandez, 2012). Hence, the most pivotal ecological processes of an ecosystem need to be identified and prioritized, in order to enable realistic decisions on which FGs are of particular concern for management actions (Mori et al., 2013).

Such a perspective on the scrutiny of biodiversity-ecosystem functioning linkages could especially bridge the gap between taxonomy- and trait-based approaches to benefit the efforts on sustaining MF that microbial diversity provides (Krause et al., 2014; Mori et al., 2013). According to the insurance hypothesis (Yachi & Loreau, 1999), an assembly of species with similar functional traits should idiosyncratically respond to external fluctuations, otherwise even slight environmental changes may give rise to the collapse of species constituting a particular functioning (Duffy, Richardson, & France, 2005; Mori et al., 2013). In light of the above, the number and the variation of soil microorganisms (*i.e.*, diversity pattern) within crucial FGs could ensure functional compensation and enable the ecosystem to maintain MF under environmental uncertainties (functional redundancy) (Elmqvist et al., 2003). Given that the diversity pattern is multidimensional and each facet has its potential distinct impact on ecosystem functioning (Bender et al., 2016; Cadotte et al., 2010; Fernando T. Maestre, Castillo-Monroy, Bowker, & Ochoa-Hueso, 2012), multiple diversity facets (*e.g.*, richness, evenness, and phylogeny-related trait dissimilarity), along with microbial abundance, are essential to sufficiently depict this ‘response-ability’ of FGs against disturbances (Bender et al., 2016; Cadotte et al., 2008; Hooper et al., 2005; Kirwan et al., 2007; Fernando T. Maestre et al., 2012; Soliveres et al., 2016). Actually, high functional redundancy of microbes has been widely assumed in natural assemblages (Bender et al., 2016; Nielsen, Ayres, Wall, & Bardgett, 2011), whilst the extent to which functional redundancy occurs is still debated (Delgado-Baquerizo, Giaramida, et al., 2016; Mori et al., 2016; Roger, Bertilsson, Langenheder, Osman, & Gamfeldt, 2016), partly due to its large variation among different FGs (Schimel & Schaeffer, 2012). With these issues in mind, it’s of practical importance to quantify the degree of multifunctional redundancy among FGs in order to disentangle the underlying mechanisms by which microbial diversity promotes MF (see Supplementary Fig. 1) (Mori et al., 2016).

To address the above concerns, we employed biological soil crusts (BSCs) as a model system (Matthew A. Bowker, Maestre, & Escolar, 2010). BSCs occur in various biomes and carry out vital EFs especially in harsh arid habitats, where severe abiotic conditions significantly constrain the growth of higher plants (Supplementary Fig. 2) (J. Belnap, 2003; M. A. Bowker, 2007). Importantly, they account for 7% of net primary production and nearly half of biological nitrogen fixation on land (J. Belnap & Lange, 2003; Elbert et al., 2012). Given the obvious carbon- and nitrogen-limited nature of arid soils (Chu et al., 2016; Dobson, Bradshaw, & Baker, 1997; Pointing & Belnap, 2012), we focused on two crucial FGs, photoautotrophic and nitrogen-fixing microorganisms. C/N-fixation by phototrophs and diazotrophs supports the heterotrophic assemblages from all domains in BSCs and further facilitates the potential establishment of higher plants (Jayne Belnap, 2002; Pointing & Belnap, 2012).

Coupling with a regional survey on the center of Tibetan Plateau, ‘the roof of our planet’ (Supplementary Fig. 3), we examined the relationships between multiple diversity facets and total abundances of phototrophs and diazotrophs and soil MF. The composition of the focal FGs was determined by using the high-throughput amplicon sequencing. Three basic α-diversity traits, as the model-based estimation of species richness, evenness, and phylogenetic dissimilarity, were quantified. In addition, seven key variables of soil functioning were measured to calculate MF, which reflect the ecological performances of BSCs on metabolic potential, soil fertility, primary productivity, carbon sequestration, and climate resistance.

We hypothesized that diversity patterns of phototrophs and diazotrophs in BSCs positively relate to MF, while the underlying linkages rely on the multiplicative impact of multiple diversity facets and ultimately depend on the extent of multifunctional redundancy in different FGs. Given the unique role of each diversity facet and their potential interactive effects (Fernando T. Maestre et al., 2012; Mori et al., 2013), we expect that multifaceted diversity explains more variance of soil MF, rather than focusing on any single ones. We further suppose that the approach used here will broaden our understanding of how an assembly of microbial species operates on MF and help to boost our practice on ecological management in drylands.

## 2. Materials and Methods

### 2.1 Study sites and sampling

The study sites are located in the largest and highest plateau on the planet, the Qinghai-Tibetan Plateau (Chen et al., 2013). We selected BSC samples from 24 sites (20 m×20 m) in different arid areas to cover a gradient of geographic and soil properties of the region (>100,000 km^2^, Supplementary Fig. 3). Most of these sites are unvegetated. Samples were taken in 2015, during the warm and wet season (May-July). Within each site, five plots (50 cm×50 cm) were selected randomly in the open field between the perennial herbs and apart from the nearest shrubs (if present). Upper BSC cores (0∼2 cm) were collected in triplicate by a 70-mm Ø ring knife in each plot, and cores from the same site were bulked and homogenized. Samples were air-dried to constant weight in a short time, packed in paper bags, and then stored in a portable refrigerator immediately. After the field survey, samples were transferred to the lab and further ground by using mortar and pestle and sieved through a 2-mm mesh to remove gravel. Subsamples were stored at −20°C until laboratory measurements.

We obtained the climate data from an online database ‘WorldClim’. Gridded climate datasets (1 km^2^ resolution) were compiled with monthly mean, minimum and maximum surface temperature and precipitation records (Hijmans, Cameron, Parra, Jones, & Jarvis, 2005). For the temperature pattern, we first conducted a principal component analysis (PCA) to reduce the dimensionality of the data of annual mean temperature, as well as mean temperatures of the wettest, warmest and coldest quarters. The primary component value (PC1) was retained to summarize local temperature patterns, which explained 69% of the total variance. The PC1 accounted for 48%, 68%, 95%, and 66% of the variance in four temperature variables, respectively. The PC1 value of the precipitation dataset was also calculated (explained 84% of variance), which accounted for 97%, 67%, and 89% of the variance in annual precipitation, precipitation seasonality and precipitation of the warmest quarter. In addition, Elevation data was collected by a handheld GPS device in the field.

### 2.2 Laboratory measurement procedures

To evaluate the abundances of target FGs, qPCR was employed to measure the absolute copy numbers of the identification genes (16S rRNA and *nif*H) amplified by group-specific primer pairs. The compositions of FGs were assessed by using a high-throughput sequencing method on an Illumina MiSeq PE300 platform with the same primer pairs (see details in Supplemental Information S2) (Wen et al., 2017). MiSeq sequencing data sets are available in the National Center for Biotechnology Information (NCBI) under the accession numbers SRR5576302 and SRR5576303.

Soil variables were measured at each site, including total nitrogen (TN), total phosphorus (TP), soil pH, dissolved ionic concentrations, total organic carbon (TOC), water-holding capacity (WHC), and chlorophyll contents (Chl) (Supplementary Tables 1 and 2). To measure soil pH, TN and TP, subsamples were first suspended in centrifuge tubes with 1M KCl (1:5 ratio, w/w), then vortexed for 1 min and shaken for 30 min at room temperature. TN and TP were determined using classic colorimetric methods (Sparks, 1996). For soil pH, after a 60 min standing of the extract, the clear supernatant was filtered through a 0.45 μm Millipore filter and measured using a pH meter (Sartorius Intec, Germany). To measure dissolved anionic and cationic concentrations (NO_3_^-^, NO_2_^-^, PO_4_^3-^, Cl^-^ and NH_4_^+^, Na^+^, Mg^2+^, Ca^2+^, respectively), subsamples were extracted with 0.5M K_2_SO_4_ (1:5 w/w) by vortexing for 5 min and then shaking for 90 min. The extracts were 10^-1^ diluted after filtering and determined on a Dionex ICS-1600 ion chromatography system (Thermo-Fisher Scientific, USA). Soil salinity was calculated as the sum of Cl^-^, Na^+^, Mg^2+^, and Ca^2+^ concentrations. To analyze TOC, subsamples were pretreated with an excess of 3M HCl to exclude the impact of inorganic carbonates and were freeze-dried overnight. The analysis was performed on a Vario TOC/TN_b_ Select analyzer (Elementar, Germany). The WHC was measured gravimetrically that 5 g of air-dried sample was thoroughly saturated with a known amount of water and then left to drain in a perforated centrifugal tube until the last drop of water had drained. The WHC was calculated as the percent weight of water absorbed per gram of soil. Chlorophyll *a* and *b* contents were calculated after extracting with 90% ethanol in dark at 4°C for 24 h and determining spectrophotometrically.

### 2.3 Estimating biodiversity and individual EFs

Three metrics of biodiversity were measured, including richness, evenness and phylogenetic dissimilarity. More specifically, the abundance coverage-based estimator (ACE) and Shannon-Wiener index (*H’*) were quantified using EstimateS v9.1.0. The ACE index was measured based on the data set of relative abundances of species, as a proxy of richness (Chao, Hwang, Chen, & Kuo, 2000). Pielou’s evenness (*J*_*sw*_) was then calculated. To account for phylogenetic dissimilarity, mean pairwise phylogenetic distance (MPD) between all pairs of phototrophic or diazotrophic species were calculated, respectively (see details in Supplemental Information S2). In brief, the consensus phylogenetic trees incorporated with relative abundance data of species were used to compute abundance-weighted MPD value of each site. The calculation was performed in the *R* package ‘Picante’ with 999 randomizations and 1000 iterations of null models to obtain a standardized effect size (SES) of MPD (Kembel et al., 2010). In addition, PCA was applied on ACE, *J*_*sw*_, and MPD to gain a synthetic standardized index ‘MultiDiver’, which represented the pattern of biodiversity and explained 56% and 53% of the variance in two FGs, respectively. While evaluating the overall diversity, a noteworthy issue here is the fractional overlap of species that belong to both FGs (*e.g.*, some of *Nostoc* spp., *Scytonema* spp., etc.). This may introduce overestimating of species richness if adding the ACE indices of FGs together. To avoid this problem, we conducted this study mainly under a framework of parallel comparative analyses that different FGs were checked separately. Even when synthetic indices of overall diversity were needed in the case of comparing their explanatory power with each other (see below), they were obtained by using PCA with covariance matrix rather than simple summing or averaging approaches. We believed that the issue of species overlapping would not make our results being invalid.

We identified seven key EFs as i) belowground biomass (‘BelowBio’, nucleic acid contents per gram), ii) TN, iii) TP, iv) soil available nutrients for plants (‘AN’, standardized sum of NH_4_^+^, NO_3_^-^, NO_2_^-^, and PO_4_^3-^ concentrations), v) potential primary productivity (‘Chl’, chlorophyll contents per gram), vi) TOC, and vii) WHC. They are widely used as indicators to evaluate the primary succession of BSCs (J. Belnap & Lange, 2003; M. A. Bowker, 2007; Hu & Liu, 2003).

The aforementioned variables were normalized to a unified comparable scale ranging from 0 to1 (Min-Max Standardization) (Soliveres et al., 2016). To address multicollinearity problems, Spearman’s rank correlation coefficients (*ρ*) of all pairs of predictors were measured (Supplementary Tables 3 and 4). Those highly correlated predictors (*|ρ|*>0.60) were prevented from being invoked in the same regression model. To account for potential confounding factors, all diversity, abundance, and functioning variables were corrected for co-varying environmental variability of climate and soils (Soliveres et al., 2016). Hence, the residuals for biotic predictors and EFs (response variables) were calculated by fitting multiple regression models (standardized temperature/precipitation PC1 values, elevation, soil pH, and soil salinity, as explanatory variables). The environment-corrected residual data was retained for subsequent analyses.

### 2.4 Evaluating the relations between biodiversity and soil MF

We conducted an analysis of multiple thresholds approach in the *R* package ‘Multifunc’ to evaluate the relations between diversity and MF. It concerns the problem of arbitrary thresholds and fully examines the fingerprint of diversity on MF (Byrnes et al., 2014). Multifunctionality level was measured as the number of EFs that exceeded a given percentage threshold of the maximum observed value across all sites. The measurement was performed along with a continuous gradient of thresholds from 1% to 99% with intervals of 1%. Since the outliers may introduce bias on the maximum observed value of each functioning variable, we calculated it as the average of the top three sites (Bradford et al., 2014). The biodiversity-functioning relationship was evaluated by fitting a series of general linear models (GLMs) with integrated indices (Fig. 1) and individual biotic predictors (Supplementary Fig. 4), respectively. We also fitted GLMs with synthetic ACE richness of two FGs, which is the common measurement of biodiversity (Jing et al., 2015; Loreau et al., 2001; F. T. Maestre et al., 2016; Soliveres et al., 2016), to compare its explanatory power with that of MultiDiver index. As the other commonly used measurement, Shannon index was highly consistent with ACE richness and/or Pielou’s evenness in our datasets (|*ρ|*>0.60, Supplementary Table 4), thus only synthetic ACE index was compared here.

**Figure 1.**
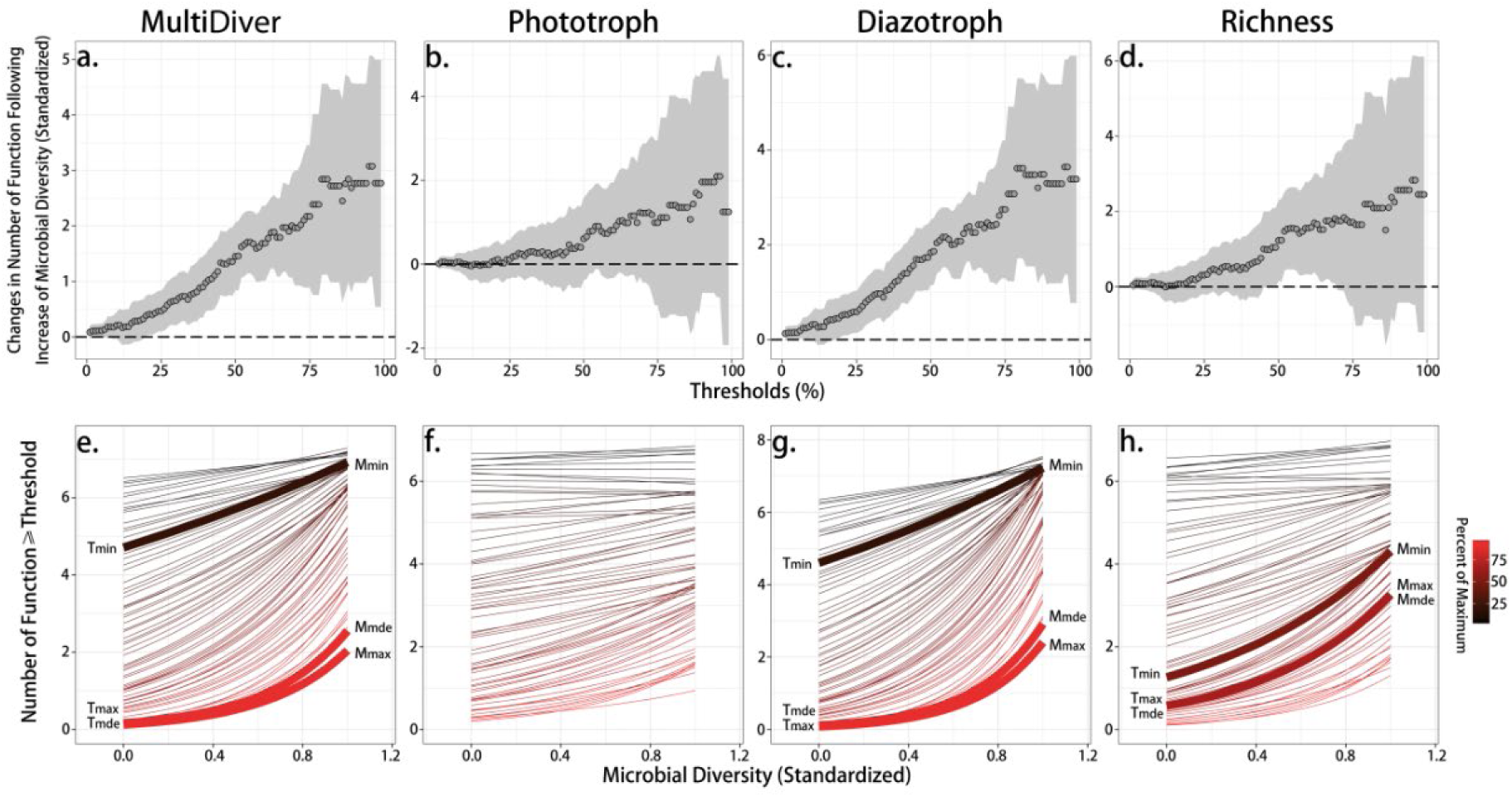
Diversity effects along with a threshold range of soil multifunctionality (MF). MultiDiver calculated by multiple metrics of diversity (*i.e.*, PC1 value of ACE richness, Pielou’s evenness, and phylogenetic dissimilarity (MPD) of both functional groups (FGs)) (a, e), synthetic phototrophic (b, f) and diazotrophic (c, g) diversity, and synthetic richness of FGs (PC1 value) (d, h) were evaluated, respectively. The upper panels (a-d) exhibited the changes in numbers of ecosystem functions (EFs) reaching a percentage threshold of the maximum observed functioning variable following one standardized unit increment of microbial diversity at multiple threshold levels. Hollow points were the fitted values of regressions and grey shading indicated 95% confidence intervals. The lower panels (e-h) showed the regression lines between diversity indices and the number of EFs above a threshold for multiple threshold values. Colors of lines indicated the thresholds from low (black) to high (red). *M*_min/max/mde_ and their corresponding thresholds *T*_min/max/mde_ were shown when the diversity effects were significantly different from zero (*i.e.*, the grey confidence interval area did not overlap the dashed line).

Three particular MF metrics were highlighted to identify key parameters (Table 1), as minimum/maximum diversity-independent MF level (*M*_min_/*M*_max_), which indicated the boundary of effects that arose due to diversity from those by chance, diversity-maximized MF level (*M*_mde_), the realized maximum effect of diversity (*R*_mde_, *i.e.*, slope), and their corresponding threshold values (*T*_min_, *T*_max_, and *T*_mde_) (Byrnes et al., 2014). To verify the possible false conclusion arising from the arbitrary selecting of EFs, we further conducted a multiple thresholds analysis on the effect of biotic predictors by removing one most highly related functioning variable with MF at each time. The result showed that the pattern of diversity effects did not change significantly (Supplementary Fig. 5). Additionally, we used an average approach to calculate soil MF level. The standardized Z-scores of seven measured EFs were averaged to obtain an MF index (F. T. Maestre et al., 2012). We then fitted linear models with environmental factors alongside uncorrected diversity predictors (Supplementary Fig. 6).

**Table 1.** Summary of multiple thresholds approach for elucidating the effects of microbial diversity on soil multifunctionality (MF).

| Predictors<sup>†</sup> | Critical Thresholds | Achieving MF levels | Estimate | Std. Error | *t*-value | Significant *P<sub>r</sub>* (>\|*t*\|) |
| --- | --- | --- | --- | --- | --- | --- |
| MultiDiver | *T*<sub>min</sub>: 0.20 | *M*<sub>min</sub>: 6.92 | 0.383 | 0.185 | 2.075 | 0.049 |
|  | *T*<sub>max</sub>: 0.99 | *M*<sub>max</sub>: 2.05 | 2.770 | 1.136 | 2.439 | 0.023 |
|  | *T*<sub>mde</sub>: 0.96 | *M*<sub>mde</sub>: 2.65 | 3.075 | 1.016 | 3.026 | 0.006 |
| Phototroph<sup>‡</sup> | - | - | - | - | - | n.s. |
| Diazotroph | *T*<sub>min</sub>: 0.19 | *M*<sub>min</sub>: 7.23 | 0.451 | 0.216 | 2.090 | 0.048 |
|  | *T*<sub>max</sub>: 0.99 | *M*<sub>max</sub>: 2.41 | 3.383 | 1.330 | 2.544 | 0.019 |
|  | *T*<sub>mde</sub>: 0.95 | *M*<sub>mde</sub>: 3.06 | 3.641 | 1.158 | 3.143 | 0.005 |
| Richness | *T*<sub>min</sub>: 0.50 | *M*<sub>min</sub>: 4.31 | 1.229 | 0.546 | 2.252 | 0.035 |
|  | *T*<sub>max</sub>: 0.67 | *M*<sub>max</sub>: 3.26 | 1.745 | 0.788 | 2.214 | 0.037 |
|  | *T*<sub>mde</sub>: 0.66 | *M*<sub>mde</sub>: 3.26 | 1.745 | 0.788 | 2.214 | 0.037 |

<sup>†</sup> synthetic single indices were calculated as the primary component values (PC1) by PCA, and the statistical parameters were obtained from the general linear models (GLMs), see details in Fig. 1; <sup>‡</sup> under the criterion of 0.95 confidence interval, phototrophic diversity in BSCs showed no significance on influencing MF through the whole range of thresholds; *T*<sub>min/max</sub>: the lowest or highest threshold where biodiversity affected MF significantly; *T*<sub>mde</sub>: the threshold where biodiversity had its strongest effect on MF; *M*<sub>min/max/mde</sub>: the number of functions achieving at the thresholds *T*<sub>min/max/mde</sub>.

Given the importance of identifying responses of individual EFs to supplement the quantifying of the overall response of MF, we used multiple linear models to evaluate the relationships between biotic predictors and seven individual EFs. This might supply additional information for understanding the possible trade-offs among different EFs (Allan et al., 2015). To implement it, we fitted paired models for each functioning variable that conflicting predictors with the multicollinearity problem were segregated into each model (Supplementary Table 4). In all case, the most parsimonious models were selected using a complete subset regression method in the *R* package ‘Leaps’. Every candidate combination of biotic predictors was tested and further simplified by removing all nonsignificant predictor according to *F*-ratio tests (significance *P*_r_(>*|t|*) >0.05). The standardized slopes were weighted by adjusted *R*^2^ of each paired model which made the slope values comparable between two models. Then, the incorporated slope was used to estimate the effect of each predictor on individual EFs.

Furthermore, we applied piecewise structural equation models (pSEM) to estimate the connection network and additive effects of biotic predictors on soil MF (Lefcheck & Freckleton, 2016). Regarding the high correlation between EFs (Supplementary Fig. 7), PCA was used to gain a PC1 value as a proxy of MF that accounted for 63% of total variance. This is an alternative option for pSEM, which is unable to invoke a latent variable, to handle the problem of highly relative variables. We constructed two independent sets of models for phototrophic and diazotrophic predictors, respectively. Shipley’s test of *d-*separation was used to check whether any paths were missing from *a priori* schematic diagram (Supplementary Fig. 8). AIC was also used to estimate the goodness-of-fit and to compare nested models. Standardized coefficients of each component model were reported along with the statistical parameters for the global fit test, such as Fisher’s *C* statistic, *p*-value, *R*^2^, AIC, as well as AICc (Shipley, 2013). The analyses were performed in the *R* package ‘piecewiseSEM’. The results of SEM were cross-checked with other approaches that we applied above, and different approaches gave very similar patterns (Supplementary Fig. 9).

### 2.5 Determining species functional importance on soil MF

Given the important roles of functional redundancy in the relations of biodiversity-ecosystem functioning (Mori et al., 2016; Nielsen et al., 2011), we quantified the multifunctional importance of two FGs at the species level, relying on randomization tests (Nicholas J. Gotelli, Ulrich, & Maestre, 2011). The species-specific values of multifunctional importance were calculated as the correlation coefficient between the target species’ relative abundance and the number of EFs which equaled to or exceeded a critical threshold across all sites. We used 999 times of randomization to reassign observed values in different sites and repeated the null modeling for all species. The SES of multifunctional importance value was then measured based on the null model. This index quantified the extent to which an observed metric deviated from a distribution of metrics generated by a stochastic simulation (N. J. Gotelli & McCabe, 2002). If |SES_*i*_| of species *i* is greater than 1.96, the value of functional importance approximately falls into the 5% tail of a normal distribution, which means that species *i* poses significantly positive or negative influence on maintaining MF, otherwise inconsiderable than expected by chance (|SES_*i*_|<1.96). We calculated SES values in a series of thresholds of EFs from 1% to 99% with an interval of 5%. Besides, we also conducted null modeling on multifunctional importance by calculating MF values via averaging the standardized Z-scores of all EFs. We implemented the randomization tests in ‘Impact’, a ‘Fortran 95’ program. In the analyses, we excluded those species that occurred only once or twice in the meta-community of the region. The mean values of SES at multiple thresholds were evaluated by simple *t*-tests, respectively (n=21), and were then checked on the difference between FGs by a paired-samples *t*-test with a two-tailed significance criterion of 0.05. Meanwhile, the patterns of SESs derived from the matrix of abundance data were compared with those by using the species presence-absence matrix. Results exhibited high consistency and therefore only former were shown here.

## 3. Results

### 3.1 Relations of microbial diversity and soil MF

As predicted, microbial diversity of overall FGs had an unambiguously strong positive effect on MF in BSCs (Fig. 1a, e). This combined influence began to be significant from a low threshold of 20% (*T*_min_), and peaked at a very high threshold of 96% (*T*_mde_) with an effect size of roughly 3.08 EFs added (*R*_mde_) between the lowest and the highest diversity levels (Table 1). As the strongest effect, overall MultiDiver achieved 44% of the maximum possible effect size on MF. However, in spite that synthetic richness of two FGs still presented a positive relation with MF (Fig. 1d, h), the impact was moderate (thresholds between 50%∼67%) and much weaker than overall MultiDiver. It could only reach 25% of the maximum possible effect. Between FGs, diversity effects varied that diazotrophic diversity was positively correlated with MF (Fig. 1c, g) while phototrophic diversity showed no significant effect (Fig. 1b, f). These results remained when conducting ordinary least squares regression and using environment-uncorrected raw data instead of residuals (Supplementary Fig. 6). When MultiDiver was disaggregated to inspect individual diversity facets, different FGs exhibited distinct patterns of effects on MF (Supplementary Fig. 4). For example, although phototrophic diversity as integrating was unrelated to MF, its MPD had a strong positive effect (*T*_min_=26%, *T*_mde_=95%, and *R*_mde_=3.37). In contrast, albeit all three diversity facets of diazotrophs were predominant factors to promote MF to a high level, its evenness and MPD just maintained MF during a moderate range of thresholds (approx. 25%∼75%) with a low effect size around 1.6∼2.0 (*R*_mde_). Diazotrophic richness was relatively more important to drive MF (*T*_min_=23%, *T*_mde_=97%, and *R*_mde_=3.05). Unexpectedly, despite that total abundance is often supposed to play a main role in biogeochemical processes (Bardgett & van der Putten, 2014; Soliveres et al., 2016; Winfree, Fox, Williams, Reilly, & Cariveau, 2015), we found moderately negative associations between microbial abundances of both FGs and soil MF (Supplementary Fig. 4).

### 3.2 Effects of diversity and abundance on individual EFs

Biotic predictors of two FGs explained a large proportion of variance in different EFs (mean adjusted *R*^2^=0.64, Fig. 2). They impacted on various EFs as strongly as climatic and ambient factors did (Supplementary Fig. 10). The most parsimonious models included 1.86±0.35 predictors across all EFs (mean±s.e.m.). The results showed that MPDs of both FGs and diazotrophic richness were the key drivers to affect various individual EFs positively (Fig. 2a). Only two EFs (TP and WHC) were negatively correlated to the increase of diazotrophic richness or phototrophic MPD, respectively (*p*<0.05). Most of the findings remained when using environmental-uncorrected raw data (Supplementary Fig. 10). Both phototrophic and diazotrophic abundances were significant positive explanatory variables for the belowground biomass. Nonetheless, diazotrophic abundance consistently showed negative relations with most EFs (*p*<0.05). Additionally, we did not detect obvious trade-offs between the effects of diversity facets and/or abundances on EFs, or between individual EFs (Supplementary Fig. 7).

**Figure 2.**
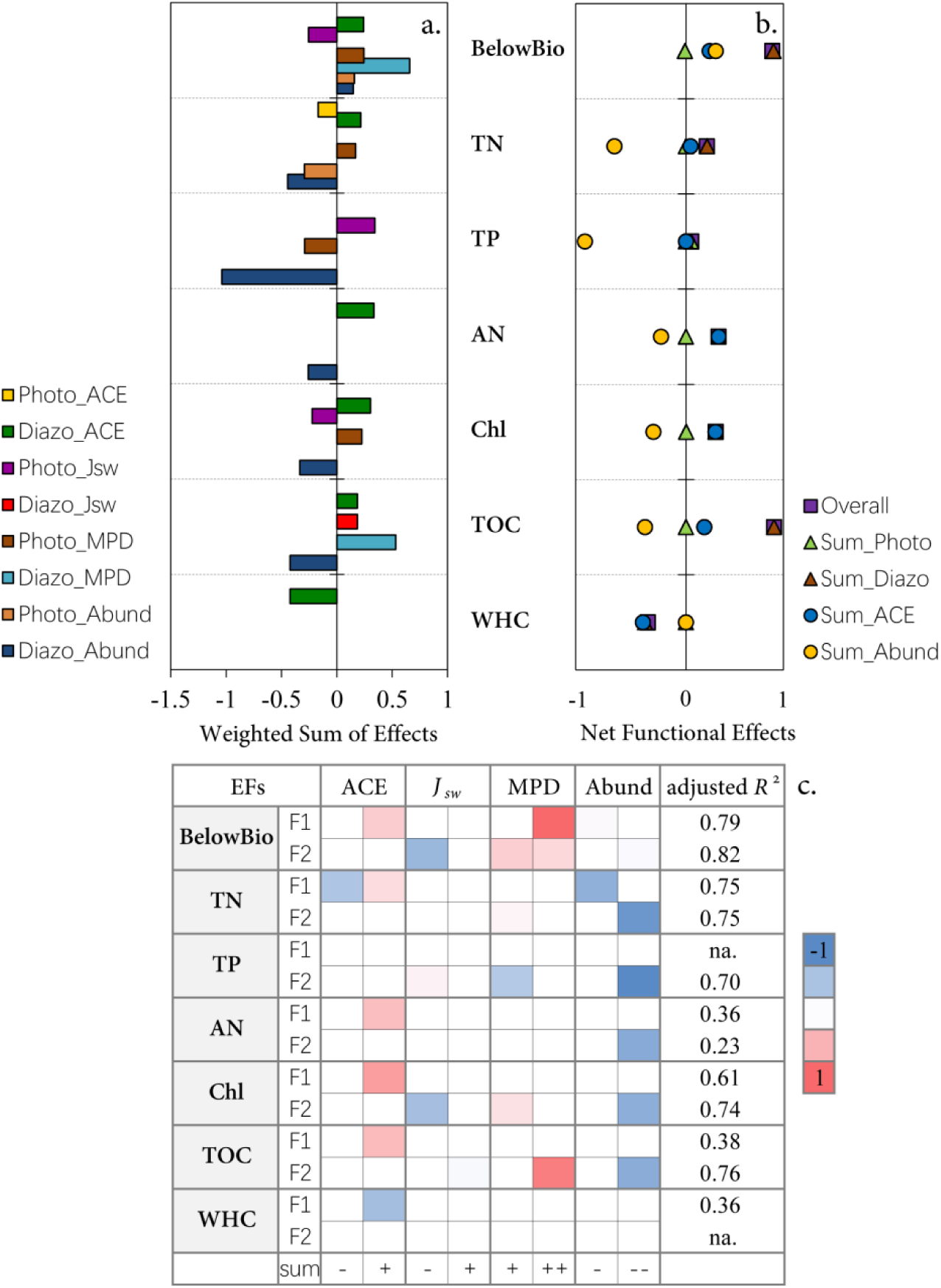
Effects of multiple biotic predictors on individual EFs. Standardized effects of diversity facets and abundances of C/N fixing FGs on each functioning variable (a), as well as the respective net effect sizes of all diversity facets (Overall), phototrophic/diazotrophic diversity facets (Sum_Photo/Sum_Diazo), and the sum of richness/abundances (Sum_ACE/Sum_Abund) were shown (b). The effect size of each predictor was the sum of estimates from the pair models (Fit 1: EF ∼ Photo_ACE + Diazo_ACE + Photo_J_sw_ + Diazo_J_sw_ + Diazo_MPD + Photo_Abund; and Fit 2: EF ∼ Photo_J_sw_ + Diazo_J_sw_ + Photo_MPD + Diazo_MPD + Diazo_Abund). Variance explained after accounting for the influence of predictor number (adjusted *R*^2^) and standardized slopes of predictors included in the pair models for each functioning variable were shown (c). Colors indicated the values of slopes from low (blue) to high (red). ‘+’ and ‘-’ indicated that the sum of the effect size of each predictor through individual EFs was positive or negative, respectively. ‘na.’ meant that no predictor had a significant effect in the most parsimonious model. ACE: ACE richness; J_sw_: Pielou’s evenness; MPD: mean pairwise distance; Abund: abundance; BelowBio: belowground biomass; TN: total nitrogen; TP: total phosphorus; AN: available nutrients; Chl: chlorophyll content; TOC: total organic carbon; WHC: water-holding capacity. ‘Photo’ and ‘Diazo’ in the figure indicated phototroph and diazotroph, respectively.

### 3.3 Connection network of diversity and abundance on soil MF

Piecewise SEM was conducted in an independent set of models for each FG, respectively (Fig. 3). Note that we did not fit the interaction of biotic predictors between two FGs, despite the fact that soil microorganisms to some extent complement with each other in supporting EFs (van der Heijden, de Bruin, Luckerhoff, van Logtestijn, & Schlaeppi, 2016). Because this would require a larger size of datasets than ours and would determine a complicated network which might be difficult to interpret. Overall, two best-fitting models had Fisher’s *C* statistic *p*>0.05 with AICc values of 53.39 and 34.85 for phototroph and diazotroph, respectively. Our SEM demonstrated that phototrophic MPD showed the highest positive direct effect on soil MF (standardized coefficient *β*=0.76). Meanwhile, it could impact MF indirectly via phototrophic richness (*β*=0.56), but phototrophic richness *per se* showed an equal negative direct effect on MF (*β*=-0.52). Besides, phototrophic evenness had a moderate negative direct influence on MF (*β*=-0.33). These direct and indirect effects of phototrophs occurred simultaneously and offset mutually. It resulted in a neutral effect of phototrophic diversity on MF as an integral factor (net coefficient *β*=-0.09), which was also observed by using multiple thresholds approach. In contrast, the relationships between phototrophic abundance and MF was weaker and nonsignificant (*β*=-0.27). For diazotrophs, only richness showed a significant positive effect on MF (*β*=0.49), and it contributed to the generally beneficial effects of diazotrophic diversity as a whole on driving soil MF.

**Figure 3.**
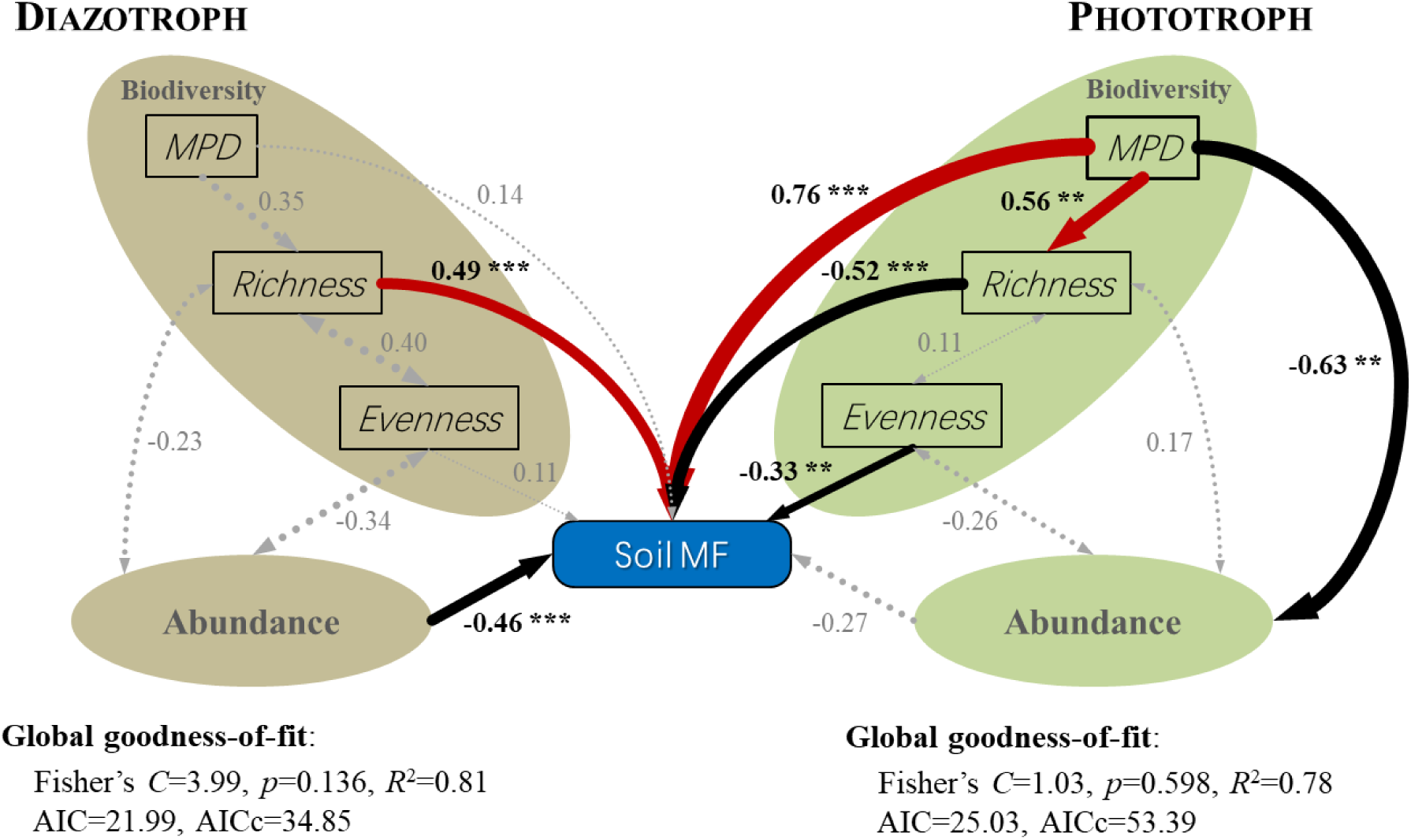
Piecewise structural equation modeling of biotic predictors (diversity and abundance) and soil MF. The connection network represented the relations among ACE richness, Pielou’s evenness, phylogenetic dissimilarity (MPD), and microbial abundances of phototrophs and diazotrophs, and their individual influences on MF. Solid red arrows indicated significant positive paths, solid black arrows indicated significant negative paths, and dotted grey arrows represented nonsignificant paths (** *p*<0.01, *** *p*<0.001). Numbers adjacent to arrows were the path standardized coefficients, and the width of arrows was proportional to the values of path coefficients. The statistical parameters for the global fit test (Fisher’s *C* statistic, *p*-value, *R*^2^, AIC, and AICc) were also shown.

### 3.4 Species functional importance on promoting soil MF

The standardized functional importance (SFI) of phototrophs showed that 7.66±0.52% (mean±s.e.m. across thresholds) of species were significantly more important to impact MF positively than the null expectation, 19.02±0.90% were negative, and another 73.32±1.23% were not functionally important (*p*>0.05) (Fig. 4a, c). The mean SFI values at thresholds from 1% to 99% were significantly lower than zero (*t*=-14.35, d*f*=20, *p*<0.001), indicating that SFIs of phototrophic species were commonly negative. For diazotrophs, 10.32±1.02% of species were significantly more important to impact MF positively than expected by chance, while only 1.77±0.64% were functionally important to impact MF negatively and were observed rarely at high threshold levels. Most diazotrophic species (87.91±1.25%) were not functionally important than the null expectation (*p*>0.05, Fig. 4b, d). Their mean SFI values at multiple thresholds were significantly higher than zero (*t*=11.27, d*f*=20, *p*<0.001), indicating that diazotrophic species generally affected MF positively. The proportion of phototrophs playing significant functional roles was over two-fold greater than that of diazotrophs, while the proportion of phototrophs with positive effect was lower. Furthermore, there was a significant difference between the SFI values of phototrophic and diazotrophic species (*t* =-12.92, d*f*=20, *p*<0.001), which implied that the extent of multifunctional redundancy between phototrophs and diazotrophs in BSCs was differentiated. In addition, the SFI calculated by averaging approach supported the findings that more proportion of diazotrophs had no significant functional importance on impacting MF, rather than phototrophs (Fig. 4e, f). That was, 159 phototrophic species functionally affected MF negatively and 50 species significantly affected MF positively (*p*<0.05), while only nine diazotrophic species posed functional importance on MF positively and 11 species were related to MF negatively (*p*<0.05).

**Figure 4.**
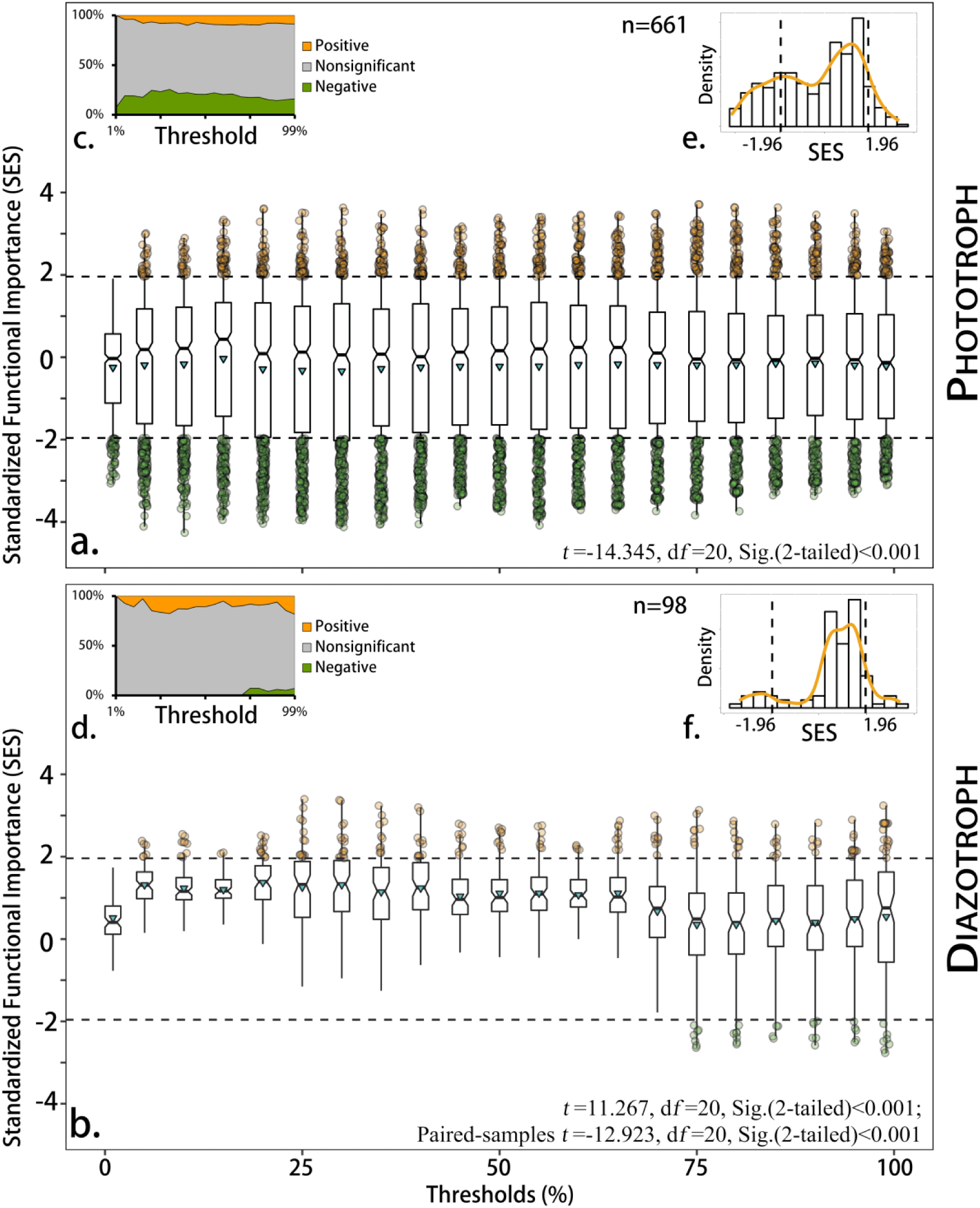
Functional importance of species to maintain multiple EFs simultaneously. Standardized functional importance (SFI) of each phototrophic (a) and diazotrophic (b) species were calculated based on multiple thresholds approach. Boxplots exhibited the distribution of SFI at every 5% threshold levels from 1% to 99%. The upper/lower quartiles, medians and average values of SFIs were shown. Black dashed lines indicated the 95% confidence intervals of SFI values. Red and green hollow points represented species with significantly positive and negative impacts on MF, respectively. Their percent proportions through the thresholds were shown (c, d). The histograms exhibited the distribution of species’ SFI values of phototrophs (e) and diazotrophs (f) based on the averaging approach. Orange lines indicated kernel density curves of SFI distributions. The statistical parameters of simple/paired-samples *t*-tests were also shown.

## 4. Discussion

### 4.1 Differentiated effects of various diversity facets on soil functionality

In our study, after separating the co-varying influence of abiotic factors, we observed a clear increase of soil MF with multifaceted diversity of the target FGs. This positive relationship appeared to reach saturation at a very high level of threshold (96%), suggesting that maintaining phototrophic and diazotrophic diversity in BSCs is important to maintain soil MF. In contrast, the synthetic richness of two FGs could just sustain soil MF during a moderate range with a relatively weak effect (50%∼67%). The result for richness here is somewhat consistent with previous studies which often claimed that, while diversity is crucial for ecosystem MF in a given considered suite of EFs, high species richness could not guarantee a ‘grand slam’ of all EFs to gain their maxima with any levels of biodiversity (Byrnes et al., 2014; Zavaleta et al., 2010). In these cases, the moderate effect of richness on MF is frequently attributed to potential trade-offs among different EFs (Byrnes et al., 2014; Mori et al., 2016). For example, in an agriculture ecosystem, increased crop yield (primary productivity) often results in declining soil formation (soil fertility) (Allan et al., 2015). An enhanced plant-available nutrient level (soil fertility), in a forest ecosystem, is very likely to associate with an increased nutrient leaching (climate resistance) (Mori et al., 2016). However, in our study, we did not detect obvious trade-offs among individual EFs (Supplementary Fig. 7). Therefore, we inferred that it’s alternatively possible that the moderate effect of species richness on MF arises from the inherent susceptibility of various EFs responding to different diversity facets. Indeed, we found that phototrophic richness has a nonsignificant effect on most individual EFs, while phototrophic MPD, as a proxy for phylogeny-related trait dissimilarity, to a large extent drives various EFs.

A previous study proposed to identify ecosystem functioning into phylogenetically and physiologically ‘narrow’ and ‘broad’ processes carried out by special constrained and a wide range of microorganisms, respectively (Schimel & Schaeffer, 2012). Regardless of the cumulative effect by richness increasing (Bender et al., 2016), if every species performs the same process identically, it doesn’t matter who is more important, and *vice versa* (Schimel & Schaeffer, 2012). In other words, the underlying factors driving different EFs might vary largely that richness *per se* is possibly crucial for some ‘broad’ EFs (*e.g.*, denitrification), but for the other ‘narrow’ EFs (*e.g.*, ammonia oxidation), the presence of certain key species (*i.e.*, composition trait) seems to be more meaningful (Bender et al., 2016; Heemsbergen et al., 2004). The results demonstrated that the multifaceted diversity indices explained more than 64% of the variance in EFs, while ACE richness accounted for less than 39% (Adjusted *R*^2^). Hence, studying diversity effects of microbes needs to incorporate multiple diversity facets to assess the combined influencing pattern, otherwise microbial diversity would be underestimated on driving MF.

### 4.2 Cascading network of biodiversity and functioning

For multiple facets of diversity, we found an apparent imbalance between the respective effects of phototroph and diazotroph on soil MF. That is, diazotrophic diversity dominates and determines the overall effect on MF, while MF is relatively independent of phototrophic diversity. However, the results of pSEM suggest that this result may partly arise from a statistical fallacy (Simpson’s paradox) whereby aggregated data likely conceal underlying links (Bradford et al., 2014). The effects of disaggregated phototrophic richness, evenness and MPD offset each other and collectively lead to an overall neutral effect. Actually, phototrophic MPD is the strongest among all biotic drivers of both FGs on MF (*β*=0.76), followed by diazotrophic richness (*β*=0.49).

Abundance-weighted MPD represents the extent of evolutionary relatedness among individuals of each FG (Cadotte et al., 2010). Evolutionary relatedness presumably reflects the integrated phenotypic difference among taxa and ultimately decides inter-/intra-species interaction (Alexandrou et al., 2015; Cadotte et al., 2010). Thus, following from the competition-relatedness hypothesis (Cahill, Kembel, Lamb, & Keddy, 2008), larger MPD indicates increased trait dissimilarity and a broader niche breadth, and gives rise to resource specialization and reduced competition within individuals of the FGs (*i.e.*, more taxa co-occurring) (Ashton, Miller, Bowman, & Suding, 2010). We indeed found that phototrophic MPD significantly promotes its richness. However, we further detected that phototrophic richness and evenness are jointly related to soil MF negatively, whilst MPD is negatively related to abundance. It means that enhanced MF is associated with the fewer phototrophic species dominant, and the overwhelming majority of the rest might be dormant or inactive with low abundances. In this context, the species pool of phototrophs with high trait dissimilarity just serve as redundancy to ensure the maxima of soil MF against the fluctuation of environmental conditions during a long-term period (namely, insurance effect), rather than by the cumulative effect of richness *per se*. Our results for species importance on MF verified that relative lower fraction of phototrophic species poses a considerable positive impact on MF at multiple thresholds than expected by chance. It suggests that the functional role of some important phototrophs is fundamental for sustaining MF (Mori et al., 2016). In fact, previous studies have proven that BSCs are often dominated by a limited suite of cyanobacterial species (*e.g., Microcoleus* spp.) (Freeman et al., 2009; Garcia-Pichel, Loza, Marusenko, Mateo, & Potrafka, 2013). The trade-off between phototrophic richness and MPD is consistent with Chesson’s framework that species coexistence is driven by the interaction of niche differences and relative fitness differences, which are two types of species differences (Chesson, 2000; Mayfield & Levine, 2010). Although larger MPD arouses greater niche difference which favors phototrophs when they drop to low densities, relative fitness difference could drive a few species to dominate. Therefore, the results exhibit that the phylogenetic relatedness among phototrophs has a strong impact on their assembling and functioning, and the strength of relative fitness difference is likely to surpass that of niche difference in this process.

In contrast, we found that diazotrophic MPD has no significant effect on its richness, abundance, as well as MF. It implies that phylogenetic relatedness among diazotrophs has little impact on their assembling and functioning. Nonetheless, another possibility is that the focal molecular marker for diazotrophs, in reality, contains little detectable phylogenetic signal on their trait dissimilarity. However, given the relatively low level of species number, the stochastic effect of diazotrophic richness is likely to provide direct benefits for MF (Bender et al., 2016). As proof, we demonstrated that, in contrast to phototrophs, the majority of diazotrophs is multi-functionally nonsignificant. It indicates a relatively low redundancy of diazotroph that the number of species, namely its richness, is more important to maintain MF (Mori et al., 2016). Our results clearly exhibited the differentiated performance of multifunctional redundancy between phototrophs and diazotrophs. It partly explains the distinctive influencing patterns of multiple diversity facets between different FGs on soil MF.

### 4.3 ‘Minority report’ of the narrow FGs on soil MF

Most importantly, it should be noted that our study sheds light on the subsets of a microbial community characterized by particular ‘narrow’ functioning (C- and N-fixation), rather than on the whole community. This may give deeper insights on exploring the mechanisms of diversity effects on MF. The within-group diversity pattern is beyond the significance of whether keystone species presence, which is previously described as a straightforward change in community composition (Wagg et al., 2014). The results imply that multifaceted diversity patterns within crucial FGs likely play an important role on ensuring the response variability of BSCs to stabilize MF against environmental perturbation (Elmqvist et al., 2003; Mori et al., 2013). Meanwhile, ecological processes performed by microbes are largely density-dependent (Yang et al., 2014). In view of this fact, some ecologists argue that species loss will not impair ecosystem functioning due to the expected community-wide increases in average population abundance (density compensation), according to the mass-ratio hypothesis (Gonzalez & Loreau, 2009; Grime, 1998). We could further infer that the lack of diversity effect on functioning tends to be more observable within the FG because it’s comprised of functionally similar species. However, our results of different models demonstrate that density compensation does not seem to be the ubiquitous mechanism by which MF is maintained in BSCs. The abundance-weighted multifaceted diversity exhibits stronger explanatory power on the variance of MF than abundance in C/N fixing FGs. Furthermore, our supplementary results show that certain relatively less dominant taxonomic groups in the FGs, such as Synechococcales and Rhodospirillales, contribute disproportionately more on explaining the variance in ecosystem functioning than Oscillatoriales or Nostocales, which are the most abundant orders (Supplementary Fig. 11). It provides the opposite evidence on the mass-ratio hypothesis.

## 5. Conclusions

In conclusion, our study demonstrated that multiple diversity facets are essential to be concerned when we try to disentangle the underlying linkages between biodiversity and ecosystem functioning. Given that multifaceted diversity pattern within C/N fixing FGs is a strong predictor on the variance in EFs/MF, it could be used as a proxy of the respond variability of BSCs against external stresses. The results give a view to bridge the gap between taxonomic and functional approaches in the biodiversity-ecosystem functioning studies. Since different diversity facets have idiosyncratic performance in different FGs, they determine divergent influencing patterns on soil MF. This shift is attributed to the significant differentiation of the extent of multifunctional redundancy between phototrophs and diazotrophs. Nevertheless, due to that our study focused on particular FGs in BSCs, it is desired to further verify the results under different habitats characteristics, as well as in other key FGs of distinct ecosystems, which could ultimately help to improve the practices of dryland management against global change.

## Competing Interests

The authors declare no conflict of interest.

## Acknowledgments

We are grateful for the technical support in the sample measuring from Min Wang of the Analysis and Test Center of IHB. We also thank Steven W. Wilhelm for helpful comments on an early version of this manuscript. The authors have no conflicts of interest to declare. This research is supported by the National Natural Science Foundation of China (Grant Nos. 41877419 and 31400368), the Special Program for Fundamental Works from the Ministry of Science and Technology of China (grant No. 2014FY210700), the Funds for the Internationalization of the University of Florence (grant No. Azione22015), and is assisted by the Supercomputing Center of CAS, Wuhan Branch.

