## Supplemental information for "Multifaceted diversity traits of crucial microbial groups in biological soil crusts promote soil multifunctionality"

**Supplemental Information S1:** Background information, including a conceptual flow diagram of our study, an illustration of the main ecosystem processes occurring in BSCs, and the geographic locations of research sites.

**Supplementary Figure 1** A conceptual flow diagram surrounding the issue of detecting microbial diversity-ecosystem multifunctionality relationships.

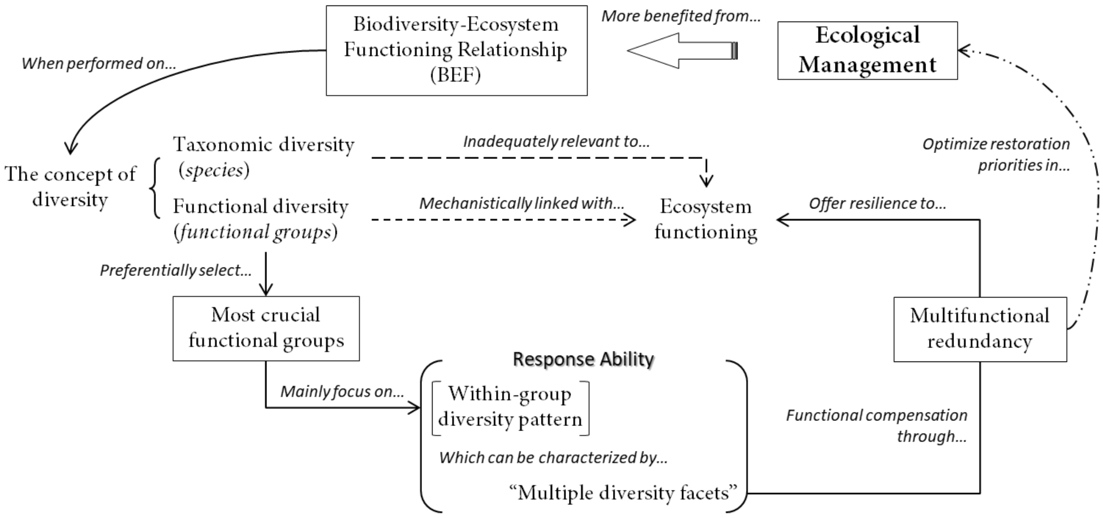

**Supplementary Figure 2** The main ecosystem processes occurring in BSCs. The biocrusts on the top layer of arid/semi-arid surfaces fulfill multiple ecological functions. This layer structure holds most bioactivities and maintains the material transfer chains and the energy flow in arid soils (after *Pointing and Belnap, 2012*, doi:10.1038/nrmicro2831).

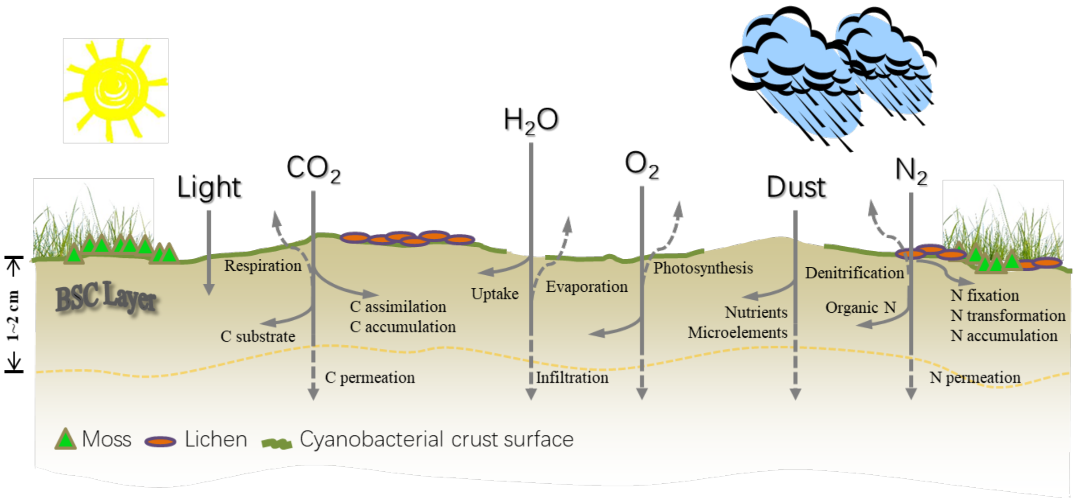

**Supplementary Figure 3** Geographic locations of sampling sites in Tibetan Plateau, "the third pole" on the earth (**a, b**). The sites were chosen for their low intensity of grazing or other land-uses in order to minimize the potential anthropogenic alteration on local microbial communities. The toponyms DR: Dangreyongcuo, SX: West Selincuo, SN: South Selincuo, CD: Cuo'E lake, MJ: Mujiucuo, ZG: Zigetangcuo, CN: Cuona lake, XC: Xiaocuo'E (**b**). The typical arid and semi-arid landscapes of the sampling sites with sparse herbaceous vegetation (**c, d**). Various widely distributing BSCs in the study region that dominant the interspace between vegetation and rocks (**e, f, g**).

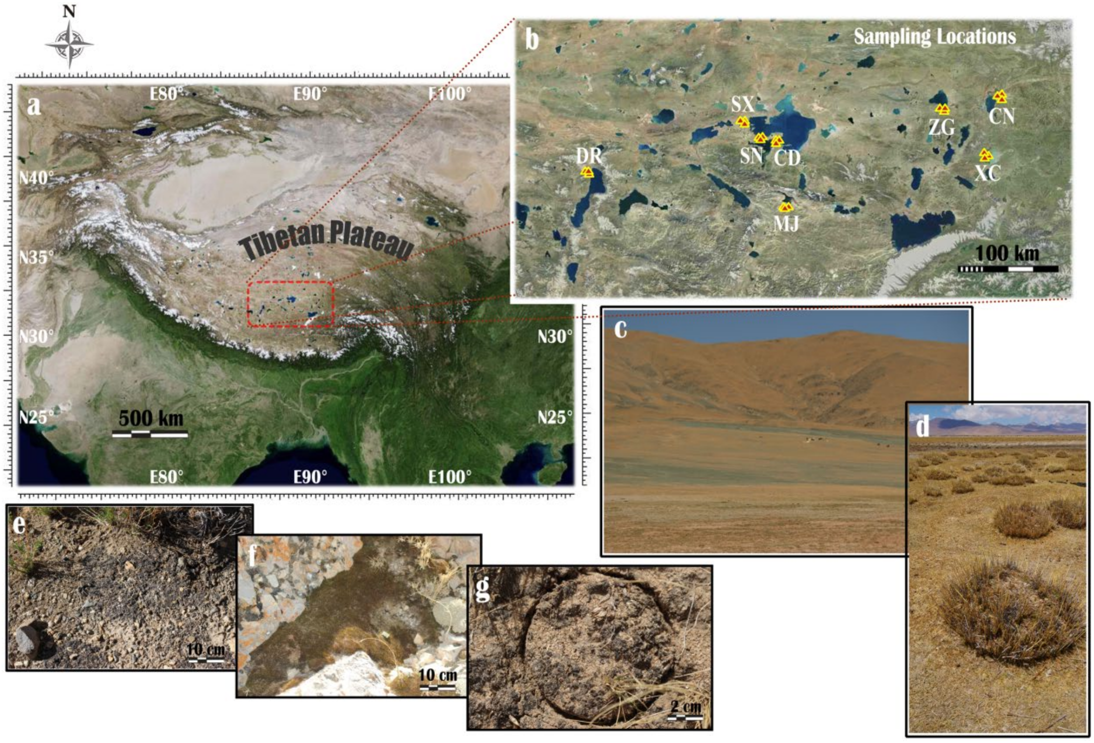

**Supplemental Information S2:** Technical details of DNA extraction, PCR amplification, high-throughput sequencing, and the clustering and annotating processes of OTUs, as well as the analysis of the linkages of dominant taxonomic groups and functioning.

### DNA extraction and qPCR

The contents of nucleic acids in the sample of each site were assessed. BSC subsamples were fully ground and mixed before the DNA extraction procedure. For each sample, 0.25 g soil was placed into a sterile 2-ml centrifuge tube. Extraction was performed following the manufacturer's protocol of a DNA isolation kit (MoBio PowerSoil DNA Kit, USA). Total DNA was eluted and dissolved in 100 μl elution buffer. DNA yields were then purified by electrophoresis with a 0.8% (w/v) agarose gel, and the bands were excised and extracted using a gel extraction kit (TaKaRa, Japan). Nucleic acid contents were quantified using a NanoDrop-8000 spectrophotometer (Thermo Scientific, USA). Extracted DNA was diluted to ~20 $ng \cdot \mu l^{-1}$ with deionized water and was stored at -80°C until subsequent molecular measurements.

To evaluate microbial abundances in BSCs, quantitative PCR (qPCR) was employed to measure the absolute copy numbers of the identification genes. For phototrophs, the group-special primer pair CYA359 (5'-GGG GAA TYT TCC GCA ATG GG-3')/CYA781 (A: 5'-GAC TAC TGG GGT ATC TAA TCC CAT T-3'; B: 5'-GAC TAC AGG GGT ATC TAA TCC CTT T-3') was used to obtain 16S rRNA gene segments of anoxygenic phototrophic bacteria, cyanobacteria and chloroplast of eukaryotic algae (Muhling, Woolven-Allen, Murrell, & Joint, 2008; Nübel, Garcia-Pichel, & Muyzer, 1997). For diazotrophs, the primer pair nifH-F (5'-AAA GGY GGW ATC GGY AAR TCC ACC AC-3')/nifH-R (5'-TTG TTS GCS GCR TAC ATS GCC ATC AT-3') was used to amplify the nitrogenase reductase gene segments (*nif*H) (Brankatschk, Towe, Kleineidam, Schloter, & Zeyer, 2011; Rosch, Mergel, & Bothe, 2002). Real-time PCR quantification assays using Power SYBR Green qPCR Master Mix (Applied Biosystems, USA) were run on an ABI-7500 Real-Time PCR

System (Applied Biosystems, USA). Each 25 μl reaction mixture contained 12.5 μl of twofold SYBR Green Master Mix, 0.5 μl of 10 μM forward and reverse primers, 2 μl of diluted DNA template, and 9.5 μl of sterile Milli-Q water. Each run contained four replicates for each sample and a blank control. The same thermal profile was performed for both functional groups, followed as an initial step at 95°C for 10 min, then 40 cycles of melting at 95°C for 15 s and annealing/elongating at 60°C for 1 min. The melting curves were obtained after each qPCR elongating step in the cycles. The copy number of target genes was interpolated from standard curves obtained from known quantities of the single-copy segment in plasmids (pMD18-T vector, TaKaRa) and run simultaneously. In detail, the plasmids, which contained inserted PCR products amplified by the primer pairs (CYA359f/781r and nifH-F/R, respectively), were $10^{-1}$ serially diluted with a $10^{-2}$ initial dilution for phototroph and $10^{-3}$ for diazotroph. For both plasmid standard curves, log-linear correlations between copies number and Ct values resulted in $R^2$>0.98. The concentrations of plasmids were determined by spectrophotometric method, and then transformed to the number of gene copies.

**High-throughput MiSeq sequencing**

Phototrophic and diazotrophic compositions in BSCs were assessed by using the high-throughput sequencing method on an Illumina MiSeq PE300 platform (Illumina Corp, USA). Briefly, the primer pairs CYA359f/781r and nifH-F/R targeting the V3-V4 hypervariable region of 16S rRNA gene and the *nif*H gene segment were synthesized with unique barcode sequences in both forward and reverse primers. The amplicon libraries were generated by PCR amplifying with TransStart FastPfu DNA polymerase (Transgen, China) and conducting on a GeneAmp-9700 PCR system (Applied Biosystems, USA). The triplicate PCR products of each sample were mixed and purified by using an AxyPrep DNA gel extraction kit (Axygen Biosciences, USA) after electrophoresis with a 2% agarose gel. Then, the extracted yields were quantified by a QuantiFluor-ST fluorometer (Promega, USA) and pooled in equimolar ratios for one MiSeq sequencing run to produce reads from both forward and reverse directions.

A total of 7.87×10$^5$ and 5.66×10$^5$ raw reads was obtained for phototrophic and diazotrophic microbes, respectively.

Raw sequencing data was processed on the Quantitative Insights Into Microbial Ecology pipeline (QIIME) incorporated with various software packages (Caporaso et al., 2010). In brief, the paired-end reads were firstly evaluated on the data quality using Trimmomatic tool v0.36 (Bolger, Lohse, & Usadel, 2014). Both forward and reverse reads were trimmed based on the average quality score of a 5-base wide sliding window if it drops below 20. Sequences without barcodes, less than 50 bases or contained undetermined "N" bases were removed. The retained sequences were sorted into the appropriate tagged samples based on their barcodes. Paired-end reads with minimum 10 bases overlap between forward and reverse read were then merged into a full length sequence by FLASH v1.2.5 (Magoc & Salzberg, 2011). The phylotypes in BSCs, namely operational taxonomic units (OTUs, referred to as species hereafter), were clustered using UPARSE tool (USEARCH v7.1) (Edgar, 2013) at 97% similarity level for phototrophic 16S rRNA gene sequences and at 95% level for diazotrophic *nif*H gene sequences (Santos et al., 2014; Wen et al., 2017) by a *de novo* packing method. A representative sequence was generated for each species by selecting the most highly connected sequence, and the taxonomic annotation of individual species was performed based on Silva Database Project (Release132, http://www.arb-silva.de) (Quast et al., 2013) and Functional Gene Repository (FunGene Release 7.3, http://fungene.cme.msu.edu/) (Fish et al., 2013). Singletons with only one sequence were deleted before the OTU packing process. To control the variation resulting from the unequal counts of sequence across samples, sequences were resampled randomly at a rarefication sequence level referred to the sample with the fewest number of sequences (Wen et al., 2017). A total of 4.29×10$^5$ and 1.65×10$^5$ filtered sequences were obtained that distributed among 873 and 146 OTUs of phototroph and diazotroph, respectively, which were used in the subsequent analyses. Cyanobacteria dominated both FGs in our research region (Supplementary Fig. 12). The sequences detected in this study caught a substantial portion of the biodiversity of both FGs in either local (α-diversity) or regional level (γ-diversity) (Supplementary

Fig. 13).

**Linkages between dominant taxonomic groups and functioning**

The dominant taxonomic groups at order level within each of two FGs were identified. We excluded the species which were not annotated to the order level from the subsequent analyses. After that, each of them had more than 5000 clean sequences across sites and consisted of species with explicit phylogenetic annotation (Supplementary Table 5). Meanwhile, those groups just comprised of species which occurred in few sites (<10 sites) were not further considered. Ultimately, ACE, $J_{sw}$, MPD, and MultiDiver index of each dominant group (six orders in phototroph and two in diazotroph) in each site were calculated accordingly. We used order-level composition data because **1)** the trait information at this taxonomic level is relatively richer, especially for cyanobacteria which is a predominant members in all sites; **2)** also, given the limit of current sequencing database particularly for functional genes, unlike higher taxonomic levels (*e.g.*, genus or class), identifying at the order-level may give more explicit annotation records (*i.e.*, less "unknown" or "unclassified").

To investigate multivariate diversity-function relationships of dominant taxonomic groups in each FG, we performed redundancy analyses that search for the liner combination of explanatory variables which explains the largest part of variance in a response matrix (individual EFs and MF). Four explanatory matrices were organized, as MultiDiver indices and abundance of the dominant taxonomic groups in each of two FGs. We used the Mantel tests (999 randomizations) to verify the significance of multivariate effects of dominant groups in phototrophs and diazotrophs, respectively. The analysis was conducted in the *R* package 'Vegan'.

On the whole, MultiDiver of dominant orders in both FGs (mean $r$=0.39) had a higher explanatory power than abundance (mean $r$=0.22) on the functioning matrix. The MultiDiver indices of Oscillatoriales, Synechococcales, eukaryotic phototrophs, and Rhodospirillales were important diversity attributes that related to Axis 1 and explained 59.79% and 20.83% of total variance, respectively ($p$<0.01, Supplementary Fig. 11a, b). Meanwhile, the abundance of Oscillatoriales, Synechococcales, and eukaryotic phototrophs also significantly correlated with a considerable variance in EFs and MF (explained 47.97% of total variance, Supplementary Fig. 11c), but the abundance of diazotrophic groups had no significant impact (Mantel test, $p$=0.148, $r$=0.17, Supplementary Fig. 11d). It's noteworthy that Synechococcales and Rhodospirillales were relatively less dominated than other groups, while they contributed more proportions on explaining the total variance of EFs and MF. The results partly contradict the mass-ratio hypothesis.

**Supplementary Figure 11** Multivariate relations between diversity and abundance of dominant taxonomic groups within each FG and the functioning matrix (EFs and MF). Redundancy analysis (RDA) displayed the linkages between dominant phototrophic (green boxes) (a, c) and diazotrophic (brown boxes) (b, d) orders and various EFs/MF variables (blue points). Solid and dashed grey arrows represented the diversity index (‘MultiDiver’) and abundance of each taxomonic group, respectively. ‘Eukaryota’ was a combined group of various eukarytic phototrophs; ‘JG30-KF-CM45’ and ‘AKYG1722’ were two orders of Thermomicrobia (Chloroflexi). The $r^2$ value of each explanatory variable was shown behind the box, and the overall degree of explantation ($r$ value) of diversity and abundance on the response matrix (EFs and MF) and the significance ($p$ value) were given by Mantel tests. The value of the axis was the variance percentage explained in the axis.

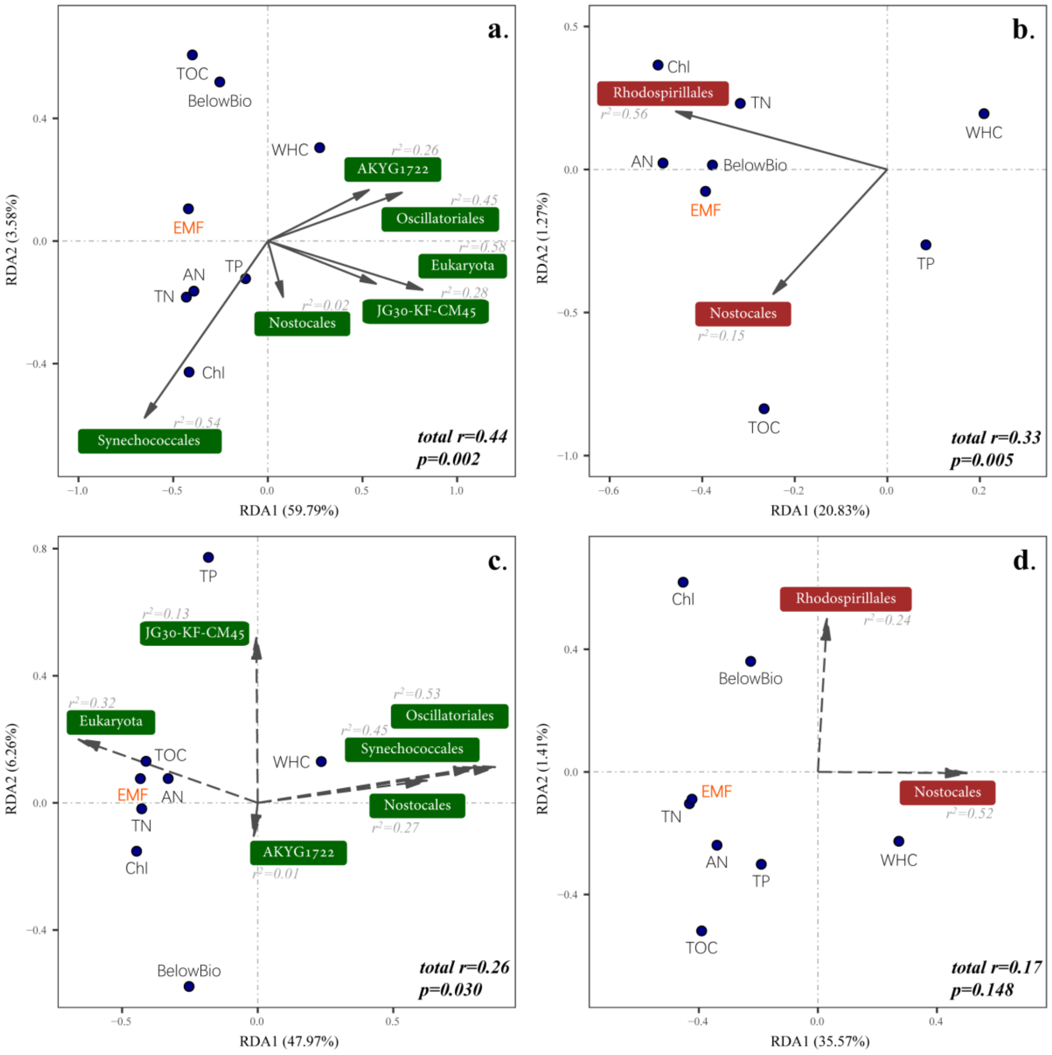

**Supplementary Table 5** Identification of dominant taxonomic groups within FGs.

| | Orders | Sequences | OTUs | Affiliation | Dominant Groups |
|---|---|---|---|---|---|
| **Phototrophs** ($4.29\times10^5$ sequences in total) | | | | | |
| **1** | Blastocatellales | 764 | 11 | Acidobacteria | |
| **2** | Acidobacteriales | 541 | 9 | Acidobacteria | |
| **3** | Armatimonadales | 130 | 10 | Armatimonadetes | |
| **4** | Herpetosiphonales | 13 | 1 | Chloroflexi | |
| **5** | Kallotenuales | 1554 | 30 | Chloroflexi | |
| **6** | Ktedonobacteria_C0119 | 76 | 4 | Chloroflexi | |
| **7** | Thermomicrobia_AKYG1722 | 8145 | 16 | Chloroflexi | √ |
| **8** | Thermomicrobia_JG30-KF-CM45 | 40545 | 151 | Chloroflexi | √ |
| **9** | Sphaerobacterales | 349 | 3 | Chloroflexi | |
| **10** | Eukaryotic phototrophs * | 99847 | 11 | Eukaryota | √ |
| **11** | Synechococcales | 25943 | 40 | Cyanobacteria | √ |
| **12** | Spirulinales | 3400 | 3 | Cyanobacteria | |
| **13** | Chroococcales | 129 | 2 | Cyanobacteria | |
| **14** | Pleurocapsales | 3495 | 2 | Cyanobacteria | |
| **15** | Chroococcidiopsidales | 2439 | 9 | Cyanobacteria | |
| **16** | Oscillatoriales | 113556 | 37 | Cyanobacteria | √ |
| **17** | Nostocales | 39323 | 15 | Cyanobacteria | √ |
| **18** | Planctomycetales | 2145 | 23 | Planctomycetes | |
| **19** | Phycisphaerales | 647 | 20 | Planctomycetes | |
| **20** | Tepidisphaerales | 1746 | 62 | Planctomycetes | |
| **21** | Chthoniobacterales | 33 | 3 | Verrucomicrobia | |
| **Diazotrophs** ($1.65\times10^5$ sequences in total) | | | | | |
| **1** | Chromatiales | 5705 | 6 | Proteobacteria | |
| **2** | Desulfuromonadales | 402 | 5 | Proteobacteria | |
| **3** | Desulfovibrionales | 2122 | 14 | Proteobacteria | |
| **4** | Rhodocyclales | 231 | 3 | Proteobacteria | |
| **5** | Burkholderiales | 6566 | 8 | Proteobacteria | |
| **6** | Rhodospirillales | 18302 | 6 | Proteobacteria | √ |
| **7** | Rhodobacterales | 1287 | 1 | Proteobacteria | |
| **8** | Rhizobiales | 99 | 2 | Proteobacteria | |
| **9** | Synechococcales | 43 | 2 | Cyanobacteria | |
| **10** | Oscillatoriales | 5356 | 3 | Cyanobacteria | |
| **11** | Nostocales | 108343 | 27 | Cyanobacteria | √ |
| **12** | Frankiales | 39 | 1 | Actinobacteria | |

* This is a combined group comprised of various eukaryotic phyla, excluding tracheophyte.

**Supplementary Figure 12** Relative phototrophic and diazotrophic abundances of the top dominant species in BSCs across all sampling sites in the region; the most dominant OTUs are sorted and organized in the genus level; the different colors represent the genus that belong to different phyla; cyanobacteria holds an important position, regarding to the functions of carbon and nitrogen fixation as the highest abundant phylum in BSCs in Tibetan Plateau; the morphological micrographs of typical cyanobacterial species in soil crusts are shown in the figure.

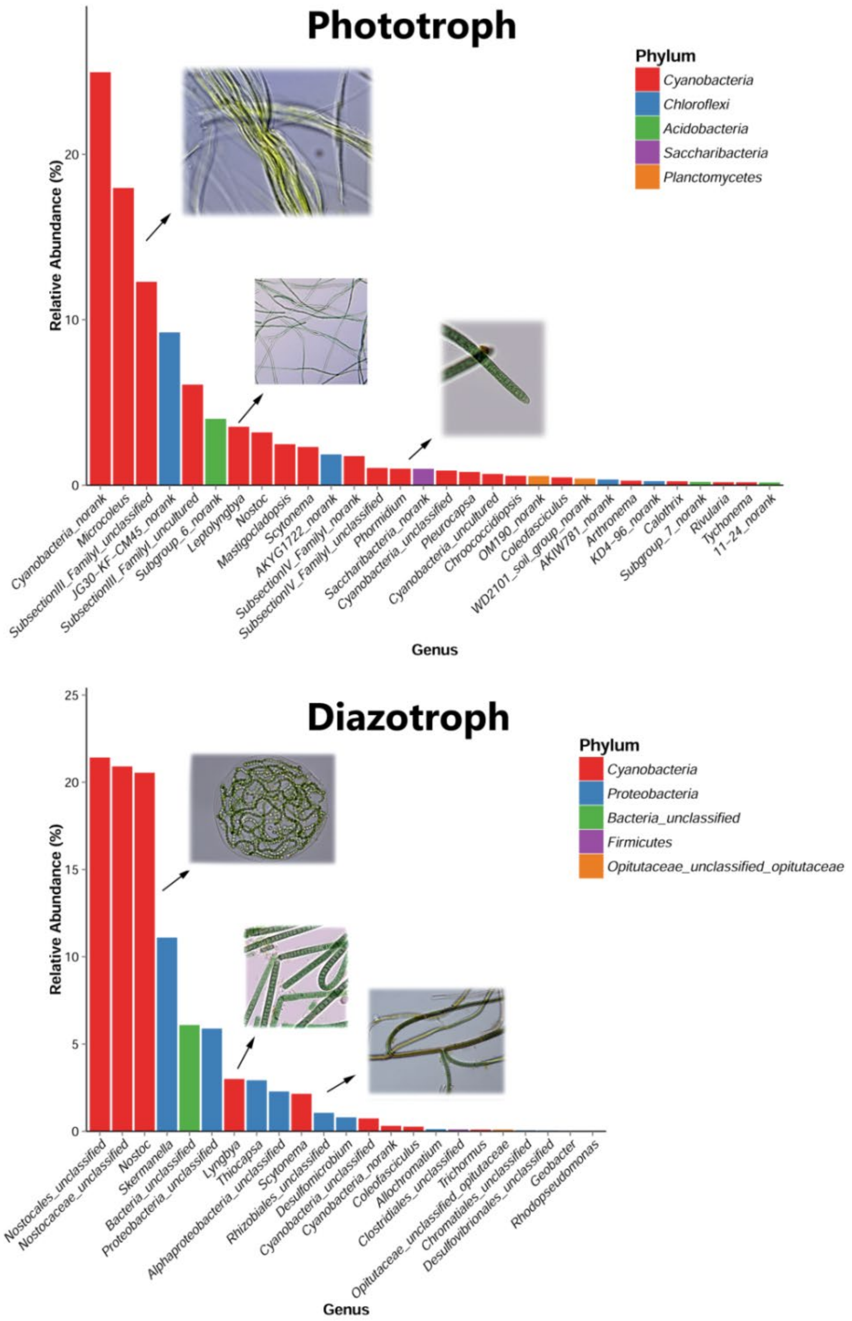

**Supplementary Figure 13** Rarefaction curves of OTUs (**a, d**) and Shannon index *H'* (**b, e**) across all sites, and species accumulation curves at the regional scale (**c, f**); the identification of phototrophic and diazotrophic OTUs are based on the similarity level of 97% and 95%, respectively; the curves demonstrate that the sequencing depth can fully describe microbial community structures in local sites as well as in the regional scale.

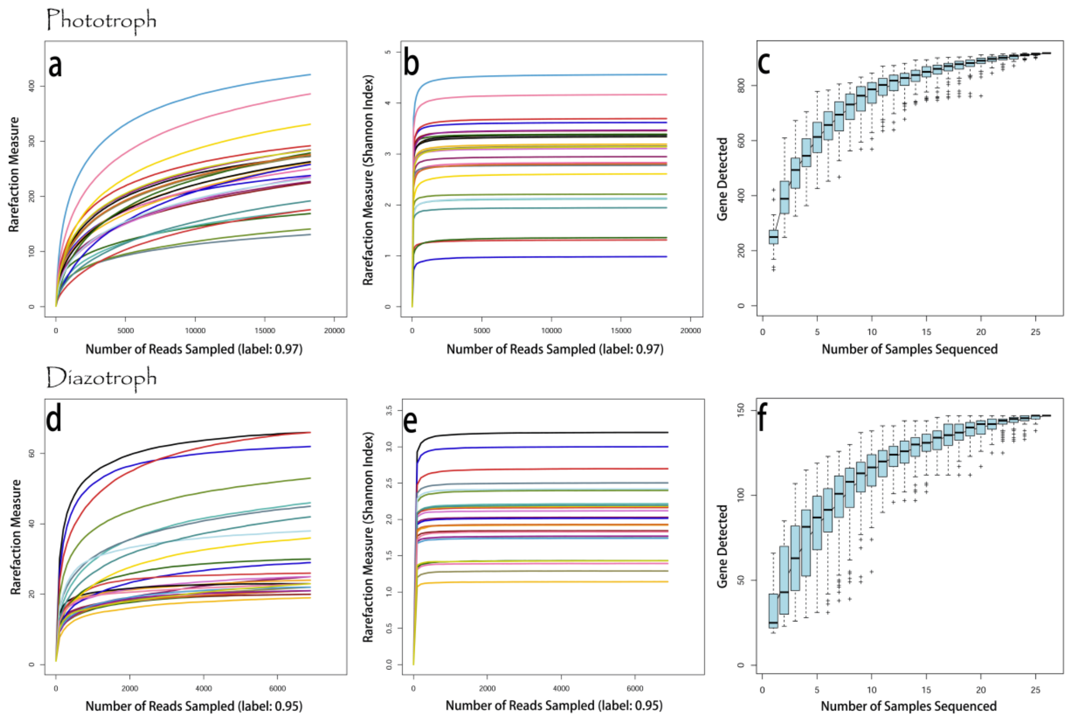

**Supplemental Information S3:** Additional statistical analyses.

**Supplementary Figure 4** Relationships between individual diversity metrics (R: ACE richness, *J*: Pielou's evenness, MPD: mean pairwise phylogenetic distance), abundances (A) and MF, evaluated by multiple threshold approach. The changes in the number of single EFs at or above a threshold of certain proportion of the maximum observed EF (open circles, thresholds 0~100%), following the increase of diversity metrics and abundances, are shown in the upper row of panels (phototroph and diazotroph, respectively), and the grey shadings indicate the 95% CI. The lines that change from black to red are from GLM regressions (set as 'quasipossion' family and 'log' link) to show the relationships between standardized R, $J_{sw}$, MPD and Aboud and the numbers of EFs at or above a threshold of some percentage of the maximum observed EF (lower row of panels); colors indicate different thresholds of maximum observed EF, and asterisks highlight the significant relationships.

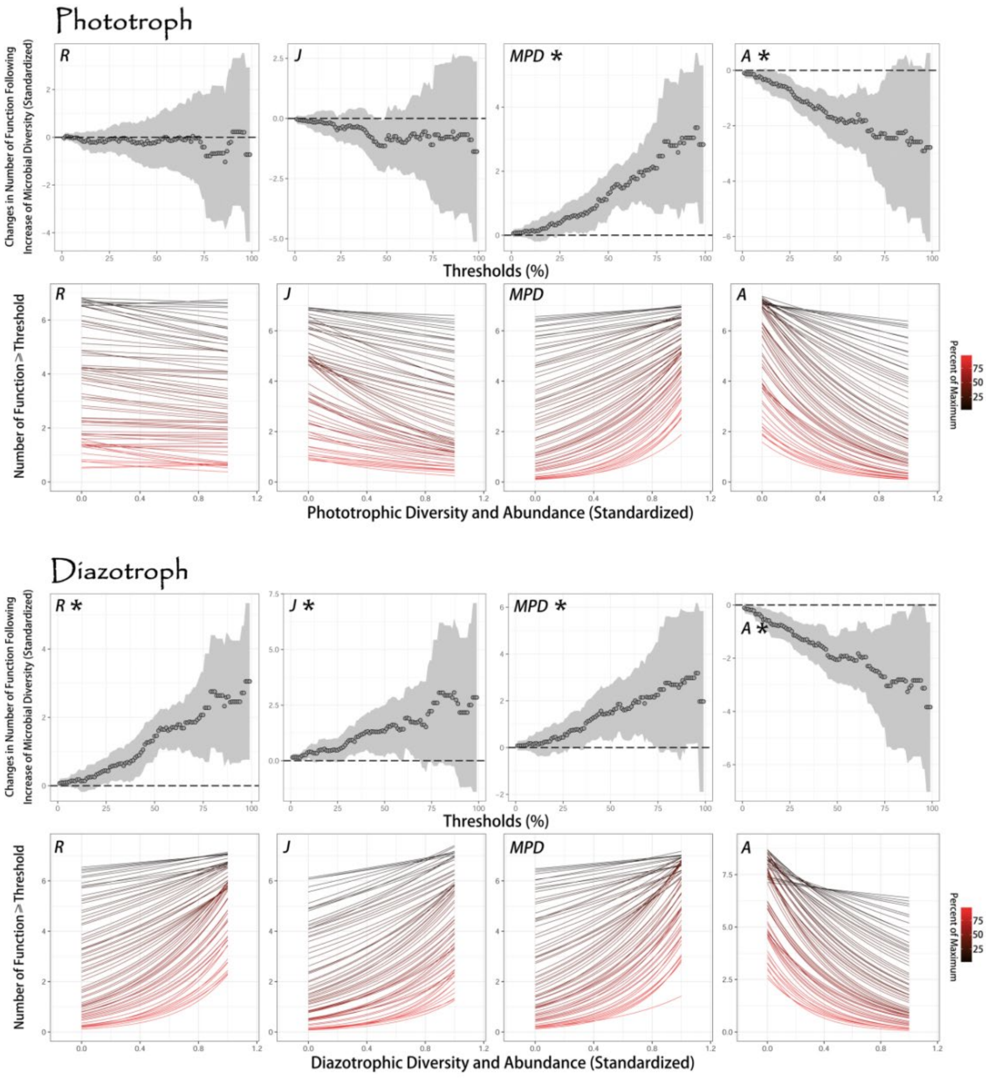

**Supplementary Figure 5** The influencing patterns of diversity and abundance on soil MF. Five different sets of data are used to estimate the relationships, as 'M' that involves all diversity predictors of both phototroph and diazotroph, 'P' that includes phototrophic diversity metrics, 'D' that includes diazotrophic diversity metrics, 'R' that only considers the richness of phototroph and diazotroph, and 'A' that involves the abundances of phototroph and diazotroph. To evaluate the potential bias of EFs that we choose, similar analyses are performed by removing one single EFs that higher correlated with multifunctionality index at each time (**a~e**), and the patterns do not change significantly.

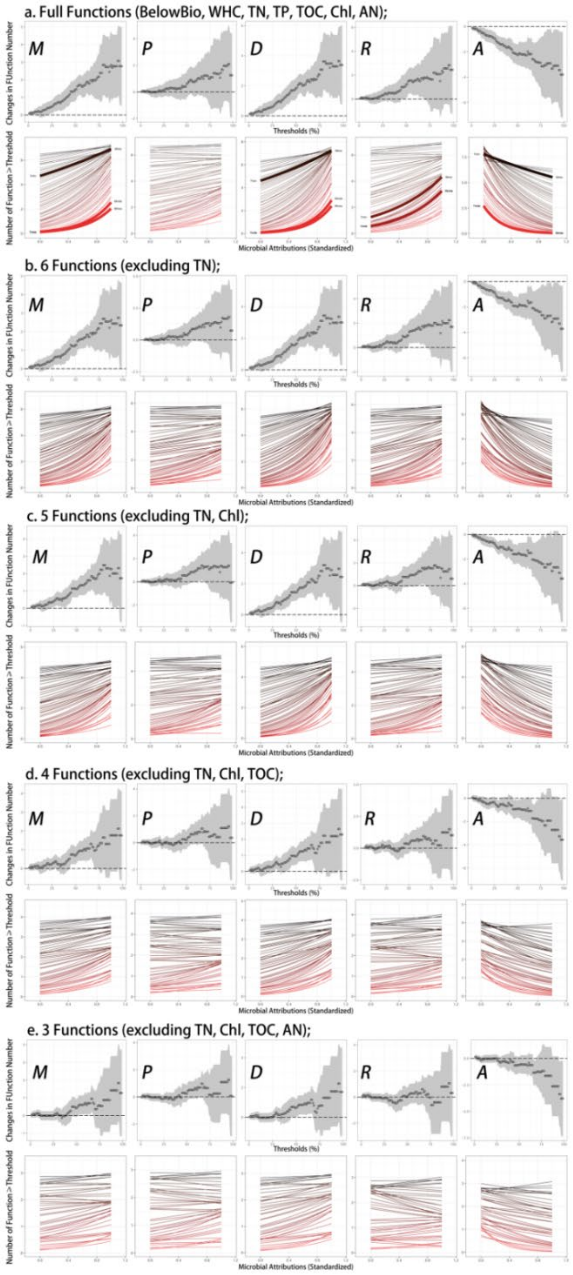

**Supplementary Figure 6** Relationships among environmental/diversity variables and soil MF. The lower triangular matrix presents the pairwise linear relationships between variables; the orange lines are fitted by using general linear models (GLM), and the dark grey areas show the 95% confidence interval; the upper triangular matrix presents Pearson correlation coefficients of each fitting; the significant level (*p* value) of the linear regressions is indicated as '***' ($p$<0.001), '**' ($p$<0.01), '*' ($p$<0.05), and '.'($p$<0.1). Temp: temperature, Prec: precipitation, a.s.l.: elevation, pH: soil pH, Salinity: soil salinity, PhotoDiv: phototrophic biodiversity, DiazoDiv: diazotrophic biodiversity, MultiFunc: MF. The data of elevation, soil pH and salinity are log-transformed. MultiFunc is the log-transformed mean value of individual EF variables. Temp, Prec, PhotoDiv, and DiazoDiv values are the principal component scores of PCA analyses.

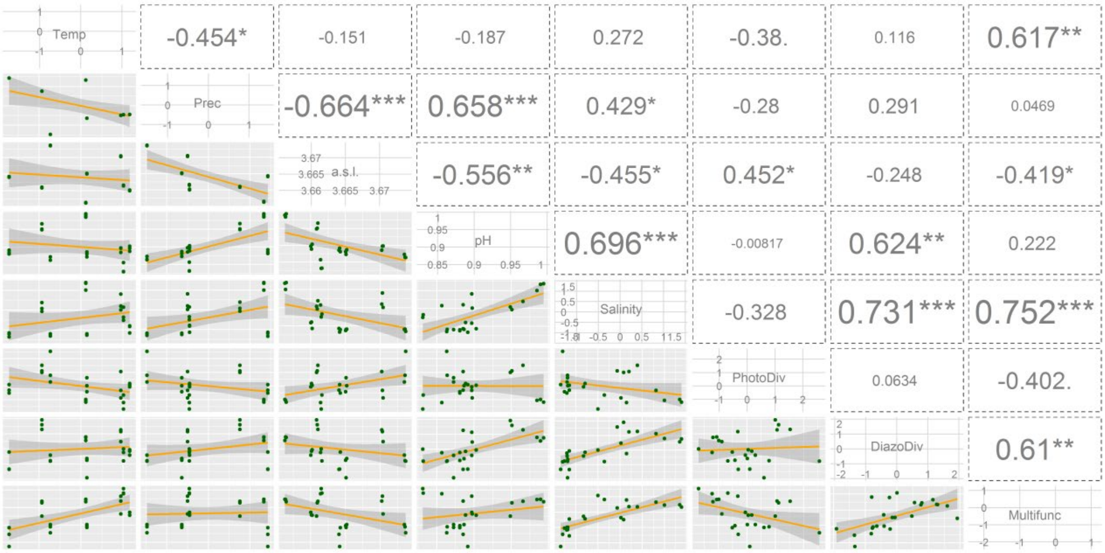

**Supplementary Figure 7** Relationships among individual EFs and MF. The lower triangular matrix presents the pairwise linear relationships between variables; the orange lines are fitted by using general linear models (GLM), and the dark grey areas show the 95% confidence interval; the upper triangular matrix presents Pearson correlation coefficients of each fitting; the significant level (*p* value) of the linear regressions is indicated as '***' (*p*<0.001), '**' (*p*<0.01), '*' (*p*<0.05), and '.' (*p*<0.1). BelowBio: belowground biomass, WHC: water-holding capacity, TN: total nitrogen content, TP: total phosphorus content, TOC: total organic carbon, Chl: chlorophyll content, AN: available nutrient, MultiFunc: MF. All variables (raw data) are log-transformed before the linear regression, and MultiFunc is the mean value of individual EFs.

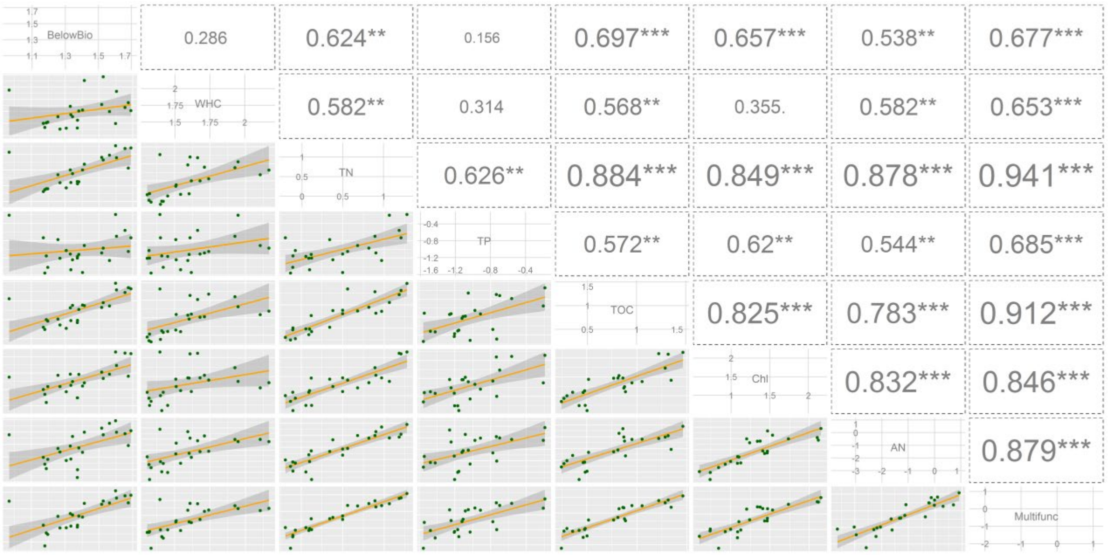

**Supplementary Figure 8** Schematic diagram describing the *a priori* influential pathways from environmental and biotic variables to soil multifunctionality (MF). Climatic characteristics are hypothesized to modulate the local soil properties like pH, salinity, etc. Environmental effects (climatic and local soil traits) are predicted to directly control the multiple functionalities of soils, and indirectly manage them through the influence on biotic components of the ecosystem. The biodiversity involves three metrics here as ACE richness, Pielou's evenness, and mean pairwise distance (MPD). Phylogenetic dissimilarity is known to influence the interspecific interaction, shape the assembling process of community, as well as correlate with the abundance (Violle et al., 2011, *doi: 10.1111/j.1461-0248.2011.01644.x*; Tan et al., 2012, *doi: 10.2307/23213510*; Peay et al., 2012, *doi: 10.1098/rspb.2011.1230*). Hence, MPD is hypothesized to directly influence the properties of species diversity (richness, evenness) and abundance (unidirectional). The richness, evenness, and abundance constitute the middle level, and each of them is driven by upper underlying predictor (MPD) and therefore appear correlated. The links among them are non-directional and interactive (double-headed arrows). Finally, all biotic predictors are predicted to directly affect the MF in soils. Paths are firstly selected based on the Spearman's correlation between the variables ($|\rho|>0.50$), and the missing paths are automatically checked by the *R* package. The latent variables are not introduced in the analysis due to the drawbacks of piecewise SEM method, despite its powerful applicability.

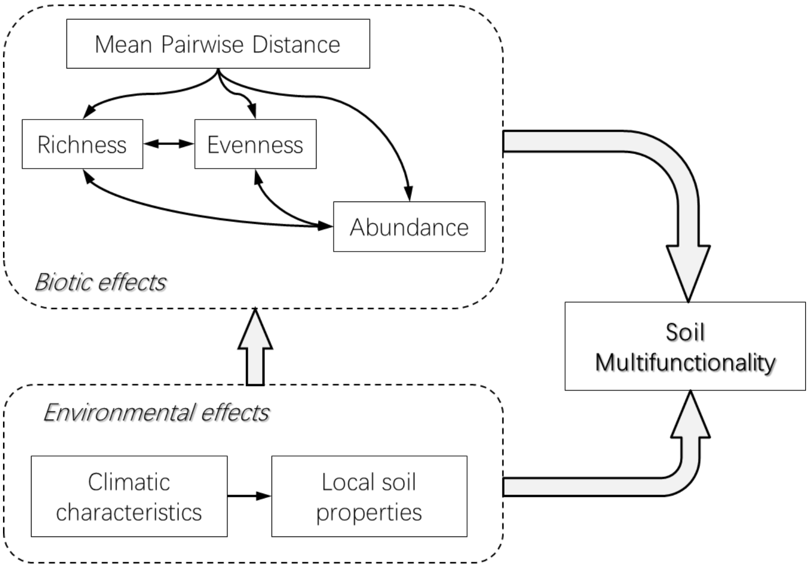

**Supplementary Figure 9** Correlations between different approaches are shown (Spearman's rank correlation coefficients $\rho$). The values in 'General models' are calculated in multiple linear regressions (ordinary least squares regression, OLS) by using standardized raw data (Raw data) or environment-corrected residual data (Residuals); the values in 'Integrated models' are the sum of standardized coefficients after weighting each one by the adjusted $R^2$ of two paired models. The correlations of values obtained from structural equation model and multiple thresholds approach are also checked, and the result demonstrates that different approaches give very similar patterns.

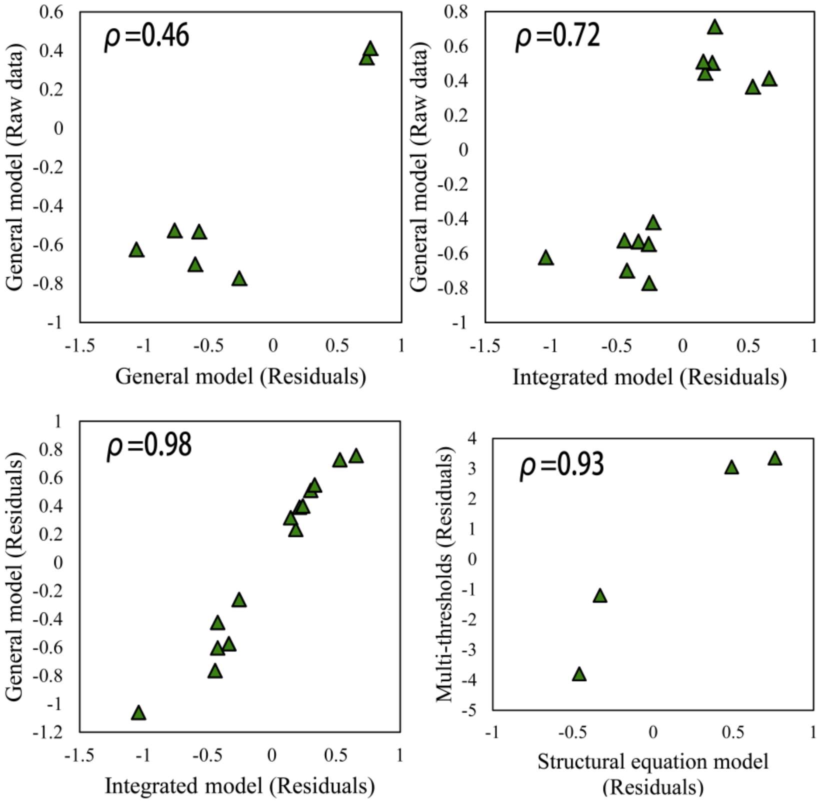

**Supplementary Figure 10** Influences of environmental and biotic factors on single EFs; standardized slope estimates (mean±s.e.m.) of significant factors ($p$<0.05) are shown, excluding the data of precipitation, pH, salinity, and diazotrophic richness and evenness due to multicollinearity problems (**a**); net functional effects of phototroph, diazotroph, and environmental factors, as the sum of related significant standardized slope estimates (**b**); the proportions of total variance in single EFs (adjusted $R^2$) that explained by multiple biotic predictors of both FGs + environmental factors (MultiDiver + Envi), phototrophic predictors + environmental factors (Photo + Envi), diazotrophic predictors + environmental factors (Diazo + Envi), and synthetic richness of two FGs + environmental factors (ACE + Envi) are shown (**c**); the comparison of four multiple regression models (Mu: MultiDiver + Envi, Ph: Photo + Envi, Di: Diazo + Envi; Ri: ACE + Envi) of predictors [from left to right, Envi (temperature, elevation, pH, and salinity), ACE (phototrophic and diazotrophic richness), $J_{sw}$ (phototrophic evenness), MPD (phototrophic and diazotrophic MPD), Abund (phototrophic and diazotrophic abundance)] and seven EFs. The color key changing from blue to red represents that the effects of predictors shift from negative to positive. Net effect sizes (sum of significant standardized slope estimates) are also shown (**d**). BelowBio: belowground biomass, TN: total nitrogen, TP: total phosphorus, AN: available nutrients, Chl: chlorophyll content, TOC: total organic carbon, WHC: water-holding capacity, Temp: temperature, Diazo: diazotroph, Photo: phototroph, Abund: abundance, ACE: ACE richness, $J_{sw}$: Pielou's evenness, MPD: mean pairwise distance.

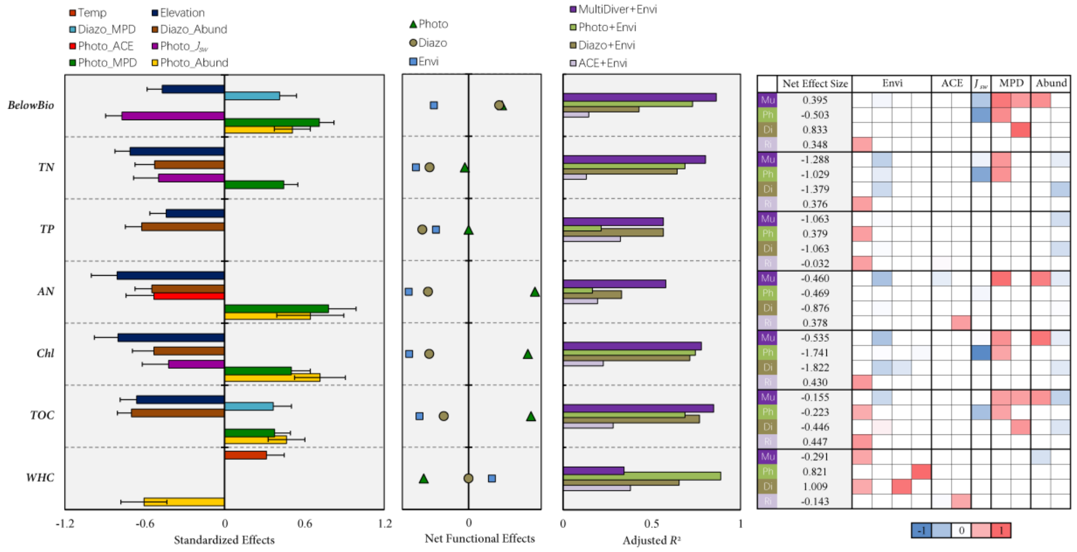

**Supplementary Table 1** Summary of climate characteristics, soil properties, biodiversity/abundance indices, and ecosystem functions used in this study.

| **Variables (*Abbr.*)** | **Description** | **Units** |
|---|---|---|
| Elevation | Geographic property. | *m* |
| Longitude | Geographic property. | *°E* |
| Latitude | Geographic property. | *°N* |
| Temperature (*Temp*) | Climatic characteristics, the principle component of the factors, including annual mean temperature, mean temperatures of wettest, warmest and coldest quarters, by PCA analysis. | *unitless* |
| Precipitation (*Prec*) | Climatic characteristics, the principle component of the factors, including annual precipitation, precipitation seasonality and precipitation of warmest quarter, by PCA analysis. | *unitless* |
| Soil pH | Soil trait, measured under 5:1 of water-soil ratio, leached by 1M KCl. | *unitless* |
| Salinity | Soil trait, sum of standardized concentrations of $Na^+$, $Cl^-$, $Mg^{2+}$, and $Ca^{2+}$. | *unitless* |
| Richness (*ACE*) | ACE richness, corrected from the number of OTUs. | *unitless* |
| Evenness ($J_{sw}$) | Pielou evenness index. | *unitless* |
| Mean Pairwise Distance (*MPD*) | Interspecies differentiation degree of each community, calculated by an integrated phylogenetic tree. | *unitless* |
| Multiple diversity index (*MultiDiver*) | Primary component of ACE, $J_{sw}$, and MPD in PCA | *unitless* |
| Abundance (*Abund*) | Copy numbers of 16S rRNA and *nifH* gene segments per gram of crust via quantitative PCR analysis. | $copies \cdot g^{-1}$ |
| Belowground Biomass (*BelowBio*) | The contents of total DNA per gram in soils, used to calculate MultiFunc. | $\mu g \cdot g^{-1}$ |
| Total Nitrogen (*TN*) | Total nitrogen contents in soils, used to calculate MultiFunc. | $mg \cdot g^{-1}$ |
| Total Phosphorus (*TP*) | Total phosphorus contents in soils, used to calculate MultiFunc. | $mg \cdot g^{-1}$ |
| Water-holding Capacity (*WHC*) | The percent of water added when soil is saturated, used to calculate MultiFunc. | *%* |
| Total Organic Carbon (*TOC*) | The percent of total organic carbon content per gram of soil (6M HCl treated), used to calculate MultiFunc. | *%* |
| Chlorophyll Contents (*Chl*) | Chlorophyll contents of crust samples (extracted by 90% alcohol), used to calculate MultiFunc. | $\mu g \cdot g^{-1}$ |
| Available Nutrients (*AN*) | Sum of standardized concentrations of $NH_4^+$, $NO_2^-$, $NO_3^-$, $PO_4^{3-}$ (leached by 0.5M $K_2SO_4$), used to calculate MultiFunc. | *unitless* |
| Multifunctionality (*MF*) | Calculated by standardized variables, as BelowBio, TN, TP, WHC, TOC, Chl, and AN. | *unitless* |

**Supplementary Table 2** Summary of climate and soil characteristics in the study sites (n=24).

| **Variables** | **min** | **max** | **range** | **mediam** | **mean** | **s.e.m.** |
|---|---|---|---|---|---|---|
| Elevation (m) | 4533 | 4716 | 183 | 4601 | 4614 | 21 |
| Longitude (°E) | 86.62 | 91.53 | 4.91 | 88.92 | 89.46 | 0.60 |
| Latitude (°N) | 31.00 | 32.02 | 1.01 | 31.68 | 31.63 | 0.12 |
| Bio1 * (℃) | 3.8 | 4.1 | 0.3 | 3.9 | 3.9 | 0.02 |
| Bio8 (℃) | 7.9 | 8.6 | 0.7 | 8.3 | 8.3 | 0.05 |
| Bio10 (℃) | 7.9 | 9.0 | 1.1 | 8.7 | 8.6 | 0.08 |
| Bio11 (℃) | -12.2 | -9.4 | 2.8 | -10.2 | -10.6 | 0.19 |
| Bio12 ($mm \cdot yr^{-1}$) | 211 | 369 | 158 | 288 | 301 | 10.5 |
| Bio15 (unitless) | 114 | 130 | 16 | 124 | 122 | 1.1 |
| Bio18 ($mm \cdot yr^{-1}$) | 149 | 257 | 108 | 214 | 215 | 6.8 |
| Soil pH (unitless) | 6.78 | 9.85 | 3.07 | 7.72 | 8.00 | 0.18 |
| $Na^+$ ($mg \cdot g^{-1}$) | 0.08 | 24.09 | 24.01 | 0.14 | 3.34 | 1.47 |
| $Ca^{2+}$ ($mg \cdot g^{-1}$) | 0.04 | 3.28 | 3.23 | 1.41 | 1.37 | 0.19 |
| $Mg^{2+}$ ($mg \cdot g^{-1}$) | 0.07 | 2.69 | 2.62 | 0.41 | 0.82 | 0.18 |
| $Cl^-$ ($mg \cdot g^{-1}$) | 0.00 | 8.73 | 8.73 | 0.02 | 0.92 | 0.45 |
| $NH_4^+$ ($mg \cdot g^{-1}$) | 0.01 | 0.35 | 0.34 | 0.06 | 0.08 | 0.02 |
| $NO_2^-$ ($mg \cdot g^{-1}$) | 0.00 | 0.34 | 0.33 | 0.01 | 0.05 | 0.01 |
| $NO_3^-$ ($mg \cdot g^{-1}$) | 0.21 | 1.80 | 1.59 | 0.35 | 0.53 | 0.09 |
| $PO_4^{3-}$ ($mg \cdot g^{-1}$) | 0.00 | 0.46 | 0.46 | 0.01 | 0.04 | 0.03 |
| TN ($mg \cdot g^{-1}$) | 0.83 | 3.58 | 2.75 | 1.56 | 1.78 | 0.18 |
| TP ($mg \cdot g^{-1}$) | 0.21 | 0.83 | 0.61 | 0.33 | 0.39 | 0.03 |
| TOC (%) | 1.76 | 28.85 | 27.09 | 6.56 | 8.42 | 1.53 |
| WHC (%) | 20.32 | 146.38 | 126.06 | 37.11 | 46.79 | 6.75 |
| Chl ($\mu g \cdot g^{-1}$) | 4.01 | 140.49 | 136.47 | 19.62 | 33.13 | 8.15 |
| Chl *a* ($\mu g \cdot g^{-1}$) | 2.25 | 88.90 | 86.64 | 13.59 | 20.48 | 4.87 |
| BelowBio DNA ($\mu g \cdot g^{-1}$) | 9.31 | 48.84 | 39.53 | 23.61 | 26.61 | 2.29 |

* The variables Bio1~18 were obtained online from WorldClim, a free database of global gridded climate datasets with a spatial resolution up to about 1 $km^2$; Bio1: annual mean temperature, Bio8: mean temperature of wettest quarter, Bio10: mean temperature of warmest quarter, Bio11: mean temperature of coldest quarter, Bio12: annual precipitation, Bio15: precipitation seasonality (coefficient of variation), Bio18: precipitation of warmest quarter.

**Supplementary Table 3** Multicollinearity check between the residuals that corrected by environmental effects.

| | Diazo_ACE | Photo_$J_{sw}$ | Diazo_$J_{sw}$ | Photo_MPD | Diazo_MPD | Photo_Abund | Diazo_Abund | Photo_*H'* | Diazo_*H'* |
|---|---|---|---|---|---|---|---|---|---|
| Photo_ACE | 0.267* | 0.058 | -0.263 | 0.627 | 0.183 | -0.348 | 0.224 | 0.330 | -0.077 |
| Diazo_ACE | | -0.389 | 0.385 | 0.711 | 0.357 | -0.526 | -0.503 | -0.239 | 0.723 |
| Photo_$J_{sw}$ | | | -0.134 | -0.155 | -0.032 | 0.091 | 0.462 | 0.905 | -0.281 |
| Diazo_$J_{sw}$ | | | | 0.209 | 0.382 | -0.305 | -0.383 | -0.185 | 0.886 |
| Photo_MPD | | | | | 0.514 | -0.612 | -0.284 | 0.082 | 0.444 |
| Diazo_MPD | | | | | | -0.530 | -0.184 | 0.081 | 0.415 |
| Photo_Abund | | | | | | | 0.649 | 0.051 | -0.459 |
| Diazo_Abund | | | | | | | | 0.550 | -0.541 |
| Photo_*H'* | | | | | | | | | -0.263 |

* Spearman's correlation between the residuals after controlling for the environmental effects; the pairs of predictors that highly correlated ($|\rho|>0.60$, shaded pink) were not invoked simultaneously in a single model; paired models, as Fit1: (multi)function~ Photo_ACE + Diazo_ACE + Photo_$J_{sw}$ + Diazo_$J_{sw}$ + Diazo_MPD + Photo_Abund, and Fit2: (multi)function~ Photo_$J_{sw}$ + Diazo_$J_{sw}$ + Photo_MPD + Diazo_MPD + Diazo_Abund, were employed to segregate conflicting predictors; Photo_ACE: phototrophic richness, Diazo_ACE: diazotrophic richness, Photo_$J_{sw}$: phototrophic evenness, Diazo_$J_{sw}$: diazotrophic evenness, Photo_MPD: phototrophic mean pairwise distance, Diazo_MPD: diazotrophic mean pairwise distance, Photo_Abund: phototrophic abundance, Diazo_Abund: diazotrophic abundance, Photo_*H'*: phototrophic Shannon index, Diazo_*H'*: diazotrophic Shannon index.

**Supplementary Table 4** Multicollinearity check between the predictors of raw data, including abiotic effects of temperature, precipitation, elevation, salinity, and soil pH.

| | Prec | Eleva | pH | Salinity | Photo_ACE | Diazo_ACE | Photo_$J_{sw}$ | Diazo_$J_{sw}$ | Photo_MPD | Diazo_MPD | Photo_Abund | Diazo_Abund | Photo_*H'* | Diazo_*H'* |
|---|---|---|---|---|---|---|---|---|---|---|---|---|---|---|
| Temp | -0.357* | -0.166 | -0.177 | 0.226 | 0.099 | -0.215 | -0.347 | 0.121 | -0.402 | 0.318 | 0.439 | -0.378 | -0.394 | -0.011 |
| Prec | | -0.709 | 0.557 | 0.310 | -0.398 | 0.578 | -0.142 | 0.126 | -0.342 | 0.189 | -0.557 | 0.307 | -0.152 | 0.311 |
| Eleva | | | -0.597 | -0.536 | 0.422 | -0.483 | 0.067 | -0.308 | 0.404 | -0.309 | 0.412 | -0.344 | 0.151 | -0.450 |
| pH | | | | 0.574 | -0.518 | 0.659 | 0.181 | 0.366 | 0.121 | 0.305 | -0.698 | 0.404 | 0.133 | 0.694 |
| Salinity | | | | | -0.421 | 0.766 | -0.407 | 0.360 | 0.051 | 0.809 | -0.480 | -0.146 | -0.461 | 0.769 |
| Photo_ACE | | | | | | -0.378 | 0.110 | -0.340 | 0.330 | -0.331 | 0.154 | -0.095 | 0.232 | -0.531 |
| Diazo_ACE | | | | | | | -0.230 | 0.335 | 0.235 | 0.693 | -0.739 | 0.041 | -0.240 | 0.780 |
| Photo_$J_{sw}$ | | | | | | | | 0.141 | 0.376 | -0.540 | -0.141 | 0.546 | 0.976 | -0.031 |
| Diazo_$J_{sw}$ | | | | | | | | | -0.046 | 0.399 | -0.246 | -0.037 | 0.098 | 0.768 |
| Photo_MPD | | | | | | | | | | 0.079 | -0.458 | 0.066 | 0.441 | 0.166 |
| Diazo_MPD | | | | | | | | | | | -0.407 | -0.468 | -0.583 | 0.696 |
| Photo_Abund | | | | | | | | | | | | -0.176 | -0.127 | -0.563 |
| Diazo_Abund | | | | | | | | | | | | | 0.556 | -0.010 |
| Photo_*H'* | | | | | | | | | | | | | | -0.089 |

* Spearman's correlation between biotic and abiotic predictors of raw data; some predictors were removed due to multicollinearity problems ($|\rho|>0.60$, shaded purple); Temp: PC1 of temperature matrix, Prec: PC1 of precipitation matrix, Photo: phototroph, Diazo: diazotroph, ACE: ACE richness, $J_{sw}$: Pielou's evenness, MPD: mean pairwise distance, Abund: abundance, *H'*: Shannon index; Temp: temperature, Prec: precipitation, Eleva: elevation.
